# Sexual dimorphism in a neuronal mechanism of spinal hyperexcitability across rodent and human models of pathological pain

**DOI:** 10.1101/2021.06.15.447407

**Authors:** Annemarie Dedek, Jian Xu, Louis-Étienne Lorenzo, Antoine G. Godin, Chaya M. Kandegedara, Geneviève Glavina, Jeffrey A. Landrigan, Paul J. Lombroso, Yves De Koninck, Eve C. Tsai, Michael E. Hildebrand

## Abstract

The prevalence and severity of many chronic pain syndromes differ across sex, and recent studies have identified differences in immune signalling within spinal nociceptive circuits as a potential mediator. Although it has been proposed that sex-specific pain mechanisms converge once they reach neurons within the superficial dorsal horn (SDH), direct investigations using rodent and human preclinical pain models have been lacking. Here, we discovered that in the Freund’s Adjuvant *in vivo* model of inflammatory pain, where both male and female rats display tactile allodynia, a pathological coupling between KCC2-dependent disinhibition and NMDA receptor potentiation within SDH neurons was observed in male but not female rats. Unlike males, the neuroimmune mediator, BDNF, failed to downregulate inhibitory signalling elements (KCC2 and STEP_61_) and upregulate excitatory elements (pFyn, GluN2B, and pGluN2B) in female rats, resulting in no effect of *ex vivo* BDNF on synaptic NMDA receptor responses in female lamina I neurons. Importantly, this sex difference in spinal pain processing was conserved from rodents to humans. As in rodents, *ex vivo* spinal treatment with BDNF downregulated markers of disinhibition and upregulated markers of facilitated excitation in SDH neurons from male but not female human organ donors. Ovariectomy in female rats recapitulated the male pathological pain neuronal phenotype, with BDNF driving a coupling between disinhibition and NMDA receptor potentiation in adult lamina I neurons following the prepubescent elimination of sex hormones in females. This discovery of sexual dimorphism in a central neuronal mechanism of chronic pain across species provides a foundational step towards a better understanding and treatment for pain in both sexes.

## Introduction

Women are disproportionately impacted by the burden of chronic pain. They are more likely than men to report low back pain, neck pain, orofacial pain, and neuropathic pain, and twice as many women report common migraines or headaches^1–3^. In quantitative evaluation of experimentally-induced pain in humans, women show more pain sensitivity than men across several noxious modalities, including mechanical-, electrical-, thermal-, and chemical-induced pain^4^. Given these epidemiological and clinical sex differences and the rising health crisis of poorly managed chronic pain, it is essential to systematically investigate the neurobiological underpinnings of pain processing in both sexes. Unfortunately, the vast majority of foundational preclinical pain research has been conducted in male rodents only^5^.

For decades it has been assumed that the molecular mechanisms that regulate neuronal excitability and behavioural outcomes within the nervous system are largely conserved between males and females. A male-biased approach has dominated neuroscience research, including pain^5,6^, typically with no rationale provided^7^. In the field of synaptic physiology, mechanisms of neuromodulation have relied almost exclusively on research in male or unsexed animals, even as examples of sexual dimorphism in synaptic circuitry and plasticity have been uncovered^8–10^. Moreover, sexual convergence onto shared phenotypic behavioural endpoints, such as pain sensitivity, may also be masking sex differences in underlying molecular and cellular mechanisms^5–7^.

Recent female-inclusive studies have revealed that pathological neuroimmune signalling in pain processing networks of the spinal dorsal horn diverges between males and females^11^. For example, the microglial TLR4/P2X4 signalling pathway that mediates spinal hyperexcitability and pain hypersensitivity in male rodent models does not have equivalent roles in female models of pathological pain^12–15^. Instead, hyperexcitability in female rodents is proposed to be driven by T-lymphocyte activation and yet-to-be-identified extracellular signals^11,13,16^. Although sex differences in immune modulation of pain have been identified, the prevailing hypothesis is that these differences converge onto conserved neuronal determinants within the dorsal horn nociceptive network^11,17^. However, specific molecular mechanisms of neuronal hyperexcitability have not been directly investigated and compared between sexes in rodent and human spinal cord.

Based primarily on evidence from male rodent models of pathological pain, a disruption in the balance between excitation and inhibition within the superficial spinal dorsal horn (SDH) is known to drive an increase in spinal nociceptive output to mediate pathological pain^18–20^. The K^+^-Cl^-^ co-transporter KCC2 plays a fundamental role in SDH inhibitory signalling by maintaining Cl^-^ gradients required for γ-aminobutyric acid receptor A (GABA_A_)- and glycine receptor (GlyR)-dependent inhibition(Kaila et al., 2014). Downregulation of KCC2, triggered by the release of brain-derived neurotrophic factor (BDNF), mediates SDH neuronal hyperexcitability in rodent models of inflammatory and neuropathic pain^12,17,22–26^. In terms of excitation at SDH synapses, BDNF also drives a potentiation of excitatory GluN2B-containing NMDARs by the Src-family kinase (SFK), Fyn, in male rodent models of pathological pain^25,27–29^. We have recently discovered that downregulation of a molecular brake, striatal-enriched protein tyrosine phosphatase-61 (STEP_61_), links BDNF/KCC2-dependent disinhibition to GluN2B NMDAR potentiation at SDH synapses in both male rodents and humans^25,27^. *In vivo* manipulation of specific molecular determinants in this pathological feed-forward mechanism has demonstrated a causative role for the above molecular targets in both SDH plasticity and pain hypersensitivity in male rodent models of chronic pain^25,26,30,31^. Whether the same neuronal determinants mediate spinal pain pathology in females remains an overlooked question of utmost clinical importance.

Here, we systematically investigated molecular mechanisms of disinhibition and NMDAR potentiation at SDH synapses across sex by measuring behavioural pain responses, changes in the distribution and abundance of synaptic proteins, and synaptic NMDAR responses in SDH neurons of males versus females. To address the translational divide between basic science rodent findings and new therapeutic approaches for humans^32^, we paired *ex vivo* BDNF and *in* vivo inflammatory rodent models of pain with a novel human spinal cord tissue preclinical model of pathological pain processing^27^. Finally, we explored whether identified sex differences in mechanisms of spinal hyperexcitability are hormonally mediated using ovariectomized female rats.

## Methods

### Animals

Rodent experiments were carried out on male or female adult (3–4-month-old) Sprague Dawley rats ordered from Charles River Laboratories. Animals were housed in same-sex pairs, on a 12-hour day/night cycle and had access to food and water *ad libitum*. Animals were randomly assigned to their respective treatment groups and were cared for in accordance with guidelines from the Canadian Council for Animal Care, Carleton University, and the University of Ottawa Heart Institute. Ovariectomized animals were ordered from Charles River Laboratories; surgery was performed at Charles River Laboratories on postnatal day 21. After animals recovered from surgery, they were shipped to the University of Ottawa Heart Institute and left to mature to age-match the intact animals.

### Freund’s adjuvant model of inflammatory pain and von Frey behaviour testing

To model persistent inflammatory pain, complete Freund’s adjuvant (CFA, Sigma) was injected into the intraplantar surface of the rats’ hindpaw. Rats were anaesthetized using isoflurane gas, and the 0.4 mL injection of either phosphate-buffered saline (PBS) vehicle or 50% v/v PBS+CFA was injected once the rat was deeply anaesthetized. Von Frey filaments were used to measure mechanical paw withdrawal threshold (PWT). PWT was measured at approximately the same time each day for all animals throughout the study. Rats were placed in testing chambers and left to acclimate for 45 minutes. PWT was measured by the simplified up-down method (SUDO)^33^ to ensure that the number of stimulus applications was standardized across subjects. Baseline testing was performed directly before vehicle/CFA injection, and was followed by testing every 24 hours until the end of the study (up to 5 days post-injection). Animals used for biochemical experiments were sacrificed 5 days post-injection, while animals used for electrophysiological recording were sacrificed 3 to 5 days post-injection. Only the ipsilateral side of the lumbar spinal cord from CFA-injected rats was used for all biochemical and electrophysiological experiments.

### Rat spinal cord isolation and preparation

Intraperitoneal injection of 3 g/kg urethane (Sigma) was used to deeply anesthetize rats. Spinal cords were then quickly dissected by ventral laminectomy and placed in ice-cold oxygenated protective sucrose solution (referred to as ‘saline’: 50 mM sucrose, 92 mM NaCl, 15 mM D-glucose, 26 mM NaHCO_3_, 5 mM KCl, 1.25 mM NaH_2_PO_4_, 0.5 mM CaCl_2_, 7 mM MgSO_4_, 1 mM kynurenic acid, bubbled with 5% CO_2_/95% O_2_). Dorsal and ventral roots were removed and the lumbar region (L3–L6) was isolated under a dissection microscope.

Spinal cords were sliced to 300 μm thickness in the parasagittal plane for electrophysiological recording using a Leica VT1200S vibratome at an amplitude of 2.75 mm and a speed of 0.1-0.2 mm/s through the dorsal horn. Slices were incubated in 34°C kynurenic acid-free saline for 40 min to promote recovery and wash out kynurenic acid. Previous control experiments have shown no difference in NMDAR synaptic responses in lamina I neurons from slices that were sectioned in saline with or without kynurenic acid, provided that slices had recovered in kynurenic acid-free saline (data not shown). Following kynurenic acid washout, the incubation chamber was removed from the heated water bath and allowed to passively cool to room temperature.

For biochemical analysis, an approximately 400 μm horizontal section containing the superficial dorsal horn was removed using a Leica VT1200S vibratome. The next 1200 μm of spinal tissue containing the deep dorsal horn and ventral horn was used for control comparisons. Following slicing, tissue for biochemical analysis was either immediately flash frozen (CFA-treated tissue) using histo-freeze (Fisher Super Friendly Freeze’It) or was treated with *ex vivo* BDNF or saline (see below) and was subsequently flash frozen.

### Human spinal cord isolation and preparation

We collected tissue from male and female adult (20–75-year-old (rounded to the nearest five for privacy)) neurological determination of death (NDD) organ donors identified by the Trillium Gift of Life Network. Candidates for donation are screened for communicable diseases (HIV/AIDS, syphilis) and conditions that could negatively affect the health of organs, such as morbid obesity. Causes of death resulted from a disruption of blood flow to/in the brain (haemorrhage or ischemia). In Canada, female deceased organ donors make up only 40% of all deceased donors^34^. Further, only 13 of the 40 total samples we have collected to date have been from female donors, resulting in a smaller sample size of female donors shown here. This difficulty in obtaining female human spinal cord tissue was one of the driving factors in why our previous study characterizing spinal determinants of pain hypersensitivity in rodents and humans^27^ included males only. Approval was obtained from the Ottawa Health Science Network Research Ethics Board for the collection of and experimentation on human spinal cord tissue. To prepare the patient for donation, hypothermia was induced using a cooling bed and the patient was perfused with a high magnesium protective solution (Celsior or Belzer UW). Following removal of organs for transplant, the spinal cord was extracted by ventral laminectomy, within 3 hours of aortic cross-clamp. Thoracic and lumbar regions were isolated and placed in ice-cold, oxygenated saline. After treatment with either BDNF or saline, tissue for western blot analysis was flash-frozen in liquid nitrogen and the dorsal horn was separated using a scalpel. Donors that had chronic pain, were taking prescription-only analgesics, or who had damaged or malformed spinal cords were excluded from western blot analysis. For immunohistochemical analysis, tissue was fixed in freshly made 4% paraformaldehyde. Donors who had damaged or malformed spinal cords were excluded from immunohistochemical analysis.

### *Ex vivo* BDNF treatment

After tissue preparation, rat or human spinal tissue was placed in oxygenated, room temperature saline containing 50-100 ng/mL recombinant BDNF (Alomone Labs) or control saline for 70 minutes. A subset of female rat experiments was carried out using an incubation time of 2-4.5 hours. Treatment of spinal tissue with BDNF and PP2 (Calbiochem) was carried out using the same approach.

### Lamina I electrophysiological recordings

Spinal cord slices were viewed under brightfield optics. Lamina I neurons were identified as being situated dorsal to the substantia gelatinosa and within 50 μm ventral to superficial white matter tracts. As previously described^25,27^, the extracellular recording solution, an artificial CSF solution, contained (in mM): 125 NaCl, 20 D-glucose, 26 NaHCO_3_, 3 KCl, 1.25 NaH_2_PO_4_, 2 CaCl_2_, and 1 MgCl_2_ in addition to 500 nM TTX, 10 μM Cd^2+^, 10 μM strychnine and 10 μM bicuculline to block voltage-gated Na^+^ channel, voltage-gated Ca^2+^ channel, glycinergic and GABAergic currents, respectively. We used borosilicate glass patch-clamp pipettes with resistances of 6–12 MΩ. The internal patch pipette solution contained (in mM): 105 Cs-gluconate, 17.5 CsCl, 10 BAPTA or 10 EGTA, 10 HEPES, 2 MgATP, 0.5 Na2GTP and had a pH of 7.25 and an osmolarity of 295 mOsm.

Neurons within lamina I were selected by size, favouring large neurons that had in increased probability of being projection neurons^35^. Whole-cell patch was established at −60 mV. Neurons were required to have an access resistance below 30 MΩ, and leakage currents less than −80 pA at a holding potential (V_h_) of −60 mV. Holding potential was slowly increased to +60 mV to record miniature excitatory postsynaptic currents (mEPSCs) that contained a dominant NMDAR-mediated slow component^36^. mEPSC traces were detected and averaged together for each treatment in Clampfit 10.7 (Molecular Devices). Selection criteria for events required that events did not completely decay within 100 ms, had an amplitude < 100 pA, and decayed to at least 50% of their overall amplitude by 500 ms. For analysis, charge transfer was measured as the area under the curve from 40-500 ms after the onset of the event. The peak amplitude of NMDAR mEPSCs was measured as the average amplitude from 18-24 ms after event onset (near the NMDAR peak, but where the AMPAR component is negligible^36^. The decay constant, τ, was derived using exponential one-term standard fitting from just after the peak of the NMDAR component to 500 ms, or where the decaying current reached steady state if earlier. For all biophysical properties of mEPSCs, measurements were taken from the average mEPSC trace for each cell and reported as an average of cells within a given treatment. After analysis, traces were transferred to Origin Pro (Northampton) for graphing.

### Western blot analysis on synaptosome fractions of rat and human spinal tissue

Synaptosomal fractions were isolated as previously described^37^. Briefly, tissue was homogenized using Wheaton dounce tissue grinders in 300 μL of ice-cold TEVP-320 mM sucrose buffer containing (in mM): 320 sucrose, 10 Tris-HCl (pH 7.4), 1 EDTA, 1 EGTA, 5 NaF, and 1 Na_3_VO_4_ with complete protease inhibitor and phosphatase inhibitor cocktails (Roche) to obtain total homogenates. The remaining total homogenate lysates were centrifuged at 4°C for 10 min at 1000g, followed by 15 min at 12 000g to obtain the crude synaptosome pellet. The pellet was resuspended in TEVP 320 mM sucrose buffer by brief sonication. The Pierce BCA protein assay kit (Thermo Scientific) was used to determine the protein content of the synaptosomal fractions. Thirty micrograms of total protein from each sample were loaded on 8% SDS-PAGE and transferred to PVDF membranes (Bio-Rad).

Membranes were blocked in 5% bovine serum albumin (BSA) in Tris-buffered saline (TBS) + 0.1% TWEEN-20 (TBS-T) and incubated overnight in 5% BSA + TBS-T plus primary antibodies [anti-STEP (1:1000), anti-KCC2 (1:1000), anti-Fyn (1:1000) and anti-β-actin (1:10 000) from Santa Cruz; anti-non-phospho-STEP (1:1000) and anti-pY416-Src (or pY420-Fyn) (1:1000) from Cell Signaling; and anti-pY1472GluN2B (1:1000) from PhosphoSolutions; anti-GluN2B (1:2000) from Millipore; for further details on antibodies used in western blots, see Supplementary Table 1. Membranes were washed three times with TBS-T and incubated in horseradish peroxidase (HRP)-conjugated secondary antibodies (anti-mouse and anti-rabbit (1:5000) from Pierce for 2 hours at room temperature. Membranes were developed using Chemiluminescent Substrate kit (Pierce) and visualized using ChemiDoc XRS+ system (Bio-Rad, Hercules, CA, USA). All densitometric bands were quantified using ImageJ (NIH).

### Immunohistochemistry

A sledge freezing Vibratome Leica VT1200S (Leica Microsystems) was used to cut 25 μm transverse sections of paraformaldehyde-fixed human spinal tissue. Sections were permeabilized in PBS (pH 7.4) with 0.2% Triton (PBS+T) for 10 min, washed twice in PBS, then incubated for 12 h at 4°C in primary anti-KCC2 antibody, anti-CGRP antibody and anti-NeuN antibody (see below) diluted in PBS+T containing 10% normal goat serum. Tissue was washed in PBS, and subsequently incubated for 2 h at room temperature in goat-Cy3 anti-rabbit purified secondary antibody (1:500, Jackson ImmunoResearch Laboratories, Cat. #111-165-144) goat anti-chicken Alexa Fluor® 647 secondary antibody (1:500, Invitrogen Cat. #AB2535866) diluted in PBS+T (pH 7.4) containing 10% normal goat serum. Spinal sections were mounted on SuperFrost™ gelatin-subbed slides (Fisherbrand) and were cover-slipped using fluorescence mounting medium (Dako, Cat. #S3023).

Nociceptive peptidergic afferent terminals, which are not present in any other types of axons in the dorsal horn^38–41^, were labelled by calcitonin gene-related peptide (CGRP) immunoreactivity using a monoclonal anti-CGRP antibody (1:5000; Sigma #C7113) raised in mouse. This antiserum detects human α-CGRP and β-CGRP but does not cross-react with any other peptide (data supplied by Sigma). We used a rabbit polyclonal antibody (1:1000, Millipore/Upstate, Cat. #07–432) to label KCC2. This antibody was raised against a His-tag fusion protein corresponding to residues 932–1043 of the rat KCC2 intracellular C-terminal^42,43^. This antibody is highly specific for rat KCC2 (KCC2a and KCC2b isoforms) and does not share any homologous sequences with other KCCs or co-transporters. We used a chicken polyclonal anti-NeuN antibody (1:1000, MilliporeSigma Cat. #6B9155 to reveal neuron cell bodies and thus to better distinguish KCC2 positive neuronal membrane.

A Zeiss LSM 880 Confocal Laser Scanning Microscope was used to acquire all confocal images. Acquisitions were 12-bit images, 2048 × 2048 pixels with a pixel dwell time of 10 μs. An oil-immersion ×63 plan-apochromatic objective was used for high magnification confocal laser scanning microscopy images, which were processed for quantification. Laser power was adequately chosen to avoid saturation and limit photobleaching. All acquisitions were performed using the same laser settings (laser, power, photomultiplier tube (PMT) settings, image size, pixel size and scanning time). The experimenter was blind to the slice treatment (saline versus BDNF treatment) and sex during acquisition.

We have previously developed MATLAB code to quantify and monitor the changes in the KCC2 intensity distributions in subcellular compartments^27^. A user must delineate the membrane of neuronal cell bodies present in the acquired confocal image. CGRP served as our marker of the SDH, and only neurons present in the regions of the SDH expressing CGRP were considered. For both imaging and analysis, the experimenter was blinded to the experimental conditions. For each pixel in the region of interest, the distance to the closest membrane segment was calculated. Using this distance map, the mean pixel intensity and standard deviation of KCC2 fluorescence signal were quantified as a function of the distance to the neuron membrane. A negative position value represents the region outside of the labelled neuron. In male humans, 143 saline-treated neurons and 205 BDNF-treated neurons from 12 adult donors were analysed. In female humans, 221 saline-treated neurons and 297 BDNF-treated neurons from 10 adult donors were analyzed. Averaged profiles were obtained for each subject and condition (saline and BDNF) and from those averages, the global KCC2 intensity profiles were obtained for each condition in each sex. The KCC2 membrane intensity (−0.5 μm < position < 0.5 μm) and the KCC2 intracellular intensity (0.6 μm < position < 2.5 μm) were extracted from each subject’s averaged KCC2 intensity profiles.

### Statistical Analysis

For all statistical analysis, p <0.05 was used as the threshold for statistical significance, and in all experiments the declared group size is the number of independent values, and statistical analysis was performed upon these values. Comparison of means for Figures 1, 2, 3E, 4, and all supplementary data were carried out using IBM SPSS Statistics 27. Parametric tests (t tests and ANOVAs) were used when assumptions of normality (tested by Shapiro-Wilk) and homogeneity of variance (tested by Levene’s Test) were met. One-way repeated-measures ANOVAs were performed to test for differences in PWT. Before running each ANOVA, we examined Mauchly’s test of sphericity. When the assumption of sphericity was violated, the Greenhouse-Geisser epsilon adjustment of degrees of freedom was used to determine the p-value. Pairwise comparisons with Bonferroni adjustment followed repeated measures ANOVAs when ANOVA achieved statistical significance (p <0.05) and showed no significant variance in homogeneity. Paired t-tests were used when comparing samples from the same rat or human (e.g. – saline versus BDNF). Prior to running paired t-tests, the normality of the data was tested (Shapiro-Wilk test), and if the data failed this test of normality (p <0.05), a Wilcoxon signed-rank test was performed instead. T-tests comparing the means from different rats were performed as independent samples t-tests. If the data failed a test of normality (Shapiro-Wilk test), then a Mann-Whitney rank sum test was performed instead. One-way ANOVAs were used when data passed assumptions of normality and homogeneity of variance and were followed by Tukey’s HSD when ANOVA achieved statistical significance (p <0.05). If the ANOVA assumption of normality was violated, the nonparametric Kruskal Wallis test was performed. If the assumption of equal variances was violated, the Welch’s test was performed, followed by Games-Howell post hoc test if Welch’s achieved statistical significance.

**Figure 1.**
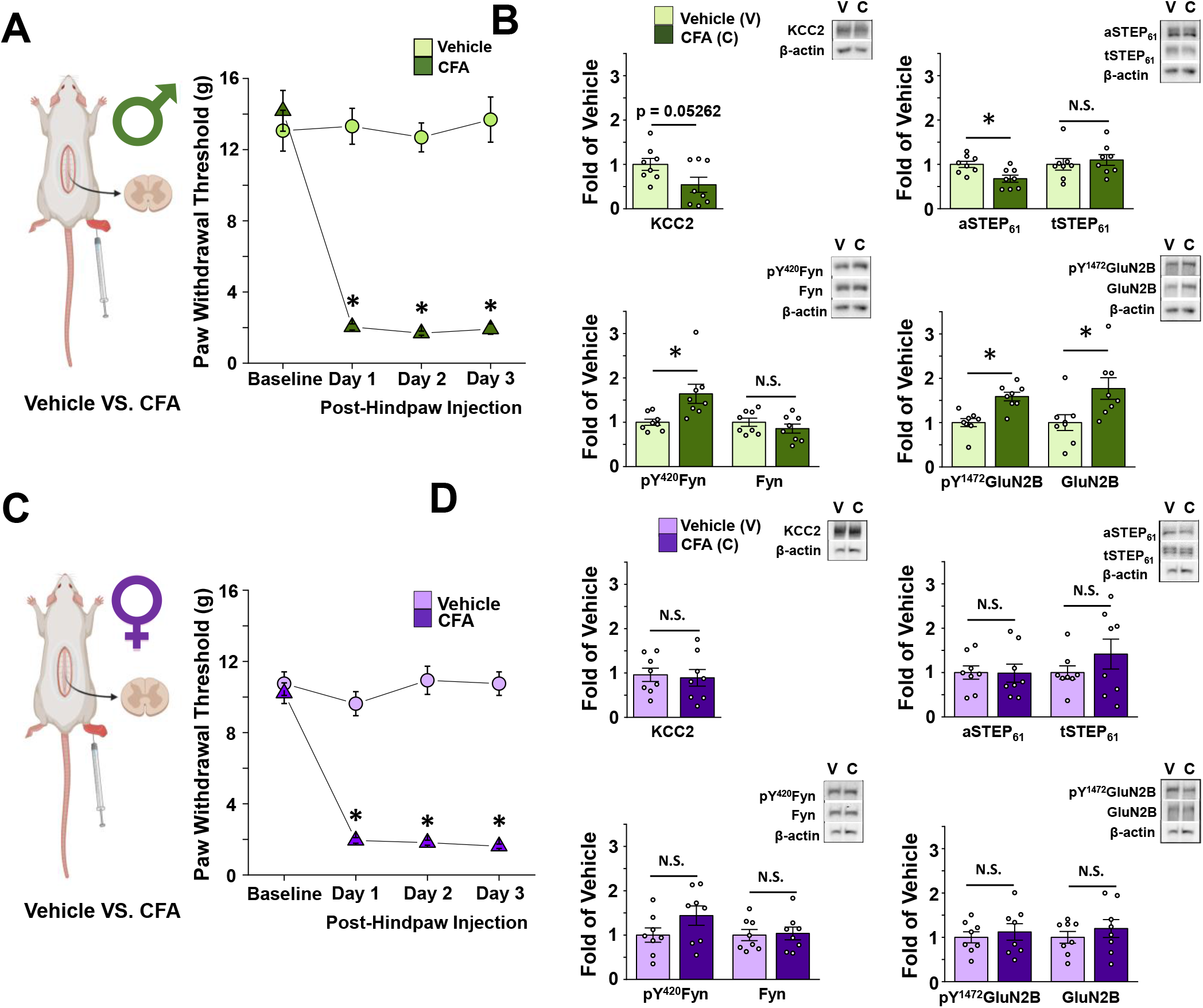
CFA elicits tactile allodynia in male and female adult rats, but only activates the STEP6i/pFyn/pGluN2B spinal hyperexcitability pathway at male SDH synapses. **A)** *Left: In vivo* CFA inflammatory pain model. *Right:* CFA elicits tactile allodynia in adult male rats. Von Frey behaviour testing shows that rats with a CFA injection (n=14) to their right hindpaw have decreased paw withdrawal threshold compared to vehicle injected animals (n=8). Animals’ tissue was collected for use in either western blot analysis or electrophysiological recordings. **B)** CFA inflammatory pain model drives upregulation of pFyn, pGluN2B, GluN2B, and downregulation of aSTEP_61_ in male rat SDH synaptosomes. Plots *(left)* and representative western blots *(right)* from SDH synaptosomes of male rats 5 days after hindpaw injection of either vehicle (light green, n=8 for all targets) or CFA (dark green, n=8 for all targets). **C)** *Left: In vivo* CFA inflammatory pain model. *Right:* Von Frey behaviour testing shows that rats with a CFA (n=14) injection to their right hindpaw have decreased paw withdrawal threshold compared to vehicle injected animals (n=8). Animals’ tissue was collected for use in either western blot analysis or electrophysiological recordings. **D)** CFA inflammatory pain model elicits no effect in the STEP_61_/pFyn/pGluN2B spinal hyperexcitability pathway of female rat SDH synaptosomes. Plots *(left)* and representative western blots *(right)* from SDH synaptosomes of female rats 5 days after injection of either vehicle (lilac, n=8) or CFA (purple, n=8). *p < 0.05

**Figure 2.**
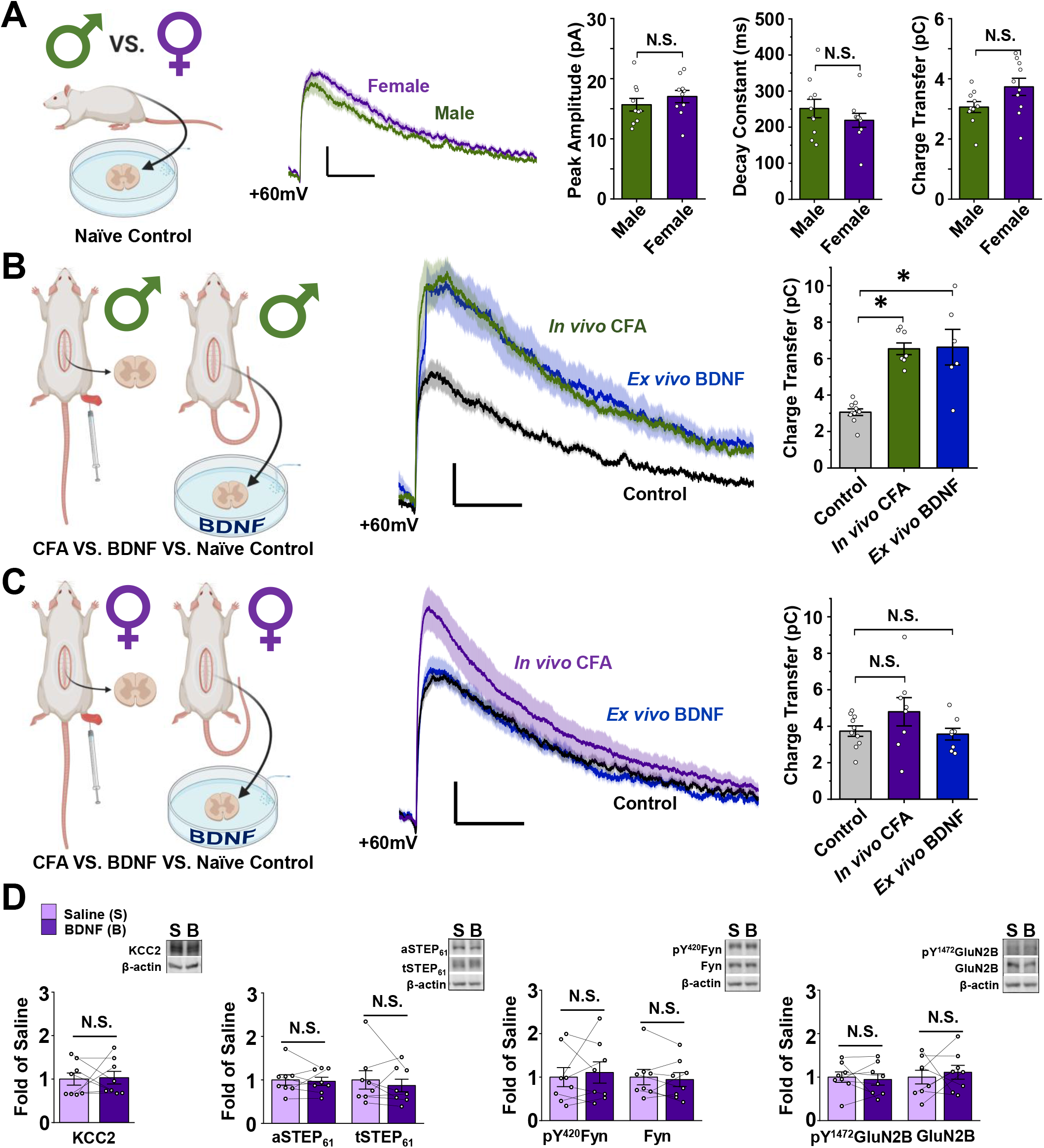
NMDARs at female SDH synapses are not potentiated or upregulated by the *ex vivo* BDNF or *in vivo* CFA models of pathological pain. **A)** Baseline lamina I mEPSCs do not differ between male and female adult SD rats. *Left:* saline treated male vs. female lamina I neurons. *Middle:* average mEPSCs at +60 mV in lamina I neurons from male (green) and female (purple) adult rats. *Right*: peak amplitude, decay constant and charge transfer of the NMDAR component of mEPSCs at +60 mV do not differ between male and female SD rats. n=10 for males and 9 for females. **B)** Male lamina I NMDAR mEPSCs are potentiated following CFA hindpaw injection and *ex vivo* BDNF treatment. *Left:* Experimental paradigm showing male *in vivo* CFA vs. *ex vivo* BDNF models. *Middle:* NMDAR mEPSCs from male rat lamina I neurons; control in black, CFA in green, BDNF in blue. *Right*, charge transfer of NMDAR mEPSCs for groups shown to left. n=10 for control, 8 for CFA, and 6 for BDNF. **C)** Female lamina I NMDAR mEPSCs are not potentiated following CFA hindpaw injection or *ex vivo* BDNF treatment. *Left*, Experimental paradigm showing female *in vivo* CFA vs. *ex vivo* BDNF models. *Middle:* NMDAR mEPSCs from female rat lamina I neurons; control in black, CFA in purple, BDNF in blue. *Right*, charge transfer of NMDAR mEPSCs shown to left. n=10 for control, 8 for CFA, and 8 for BDNF. **D)** *Ex vivo* BDNF treatment model elicits no effect in female rat SDH synaptosomes. Plots (left) and representative western blots (right) from female rat SDH synaptosomes of tissue treated with either control saline (lilac, n=8) or 50 ng/mL recombinant BDNF for 70 minutes (purple, n=8). *p < 0.05

**Figure 3.**
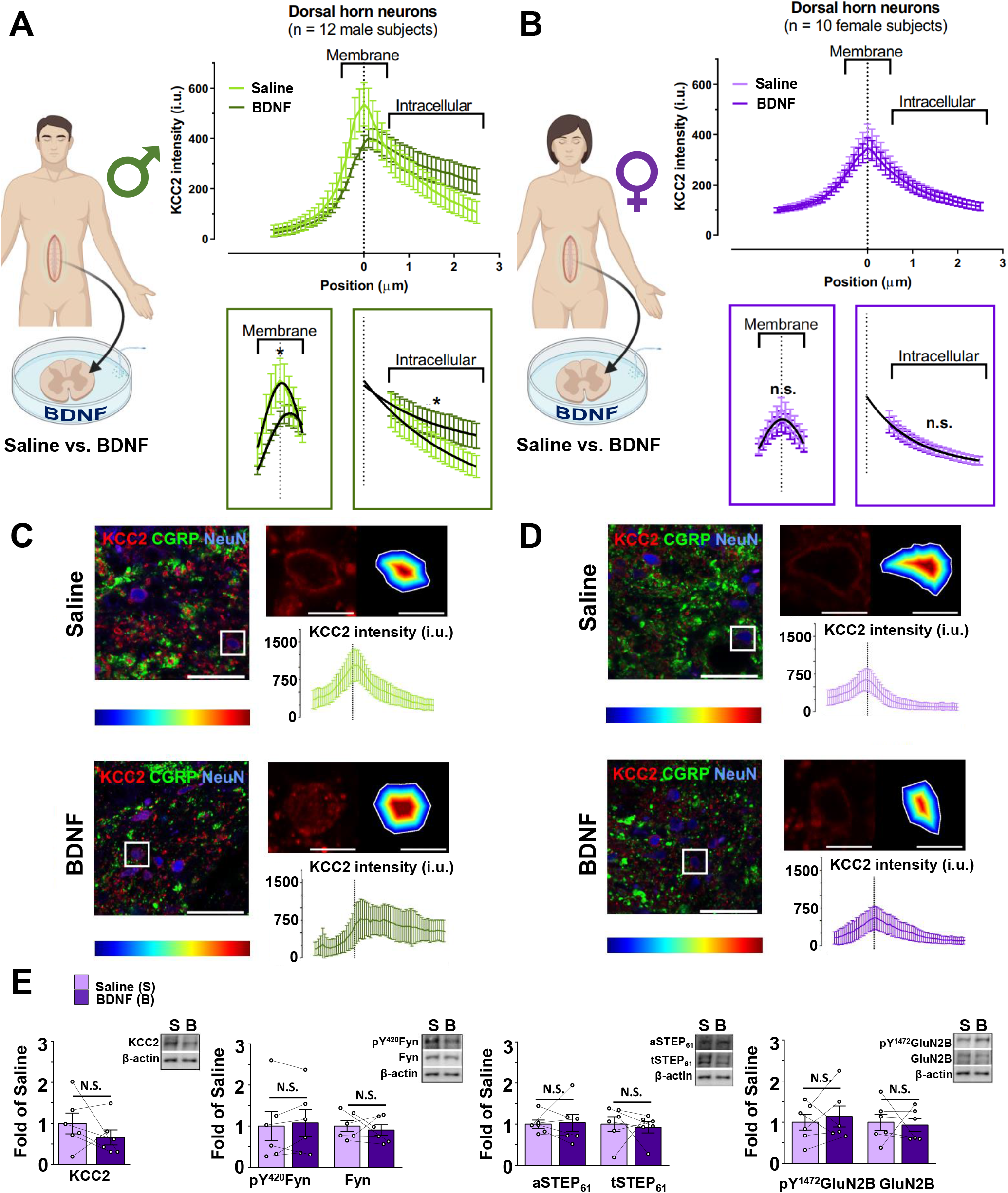
In contrast to males, *ex vivo* BDNF treatment does not activate markers of disinhibition or facilitated excitation at SDH synapses of female human spinal tissue. **A)** The *ex vivo* BDNF model elicits KCC2 internalization in adult male human SDH neurons. *Left:* experimental paradigm showing treatment of human SDH tissue in either saline or BDNF. *Right:* Average KCC2 intensity values from SDH neurons of male human donor tissue (n=12) incubated in saline (light green) versus BDNF (dark green), with comparisons of membrane and intracellular regions below. **B)** The *ex vivo* BDNF model has no effect on KCC2 internalization in adult female human SDH neurons. *Left:* experimental paradigm showing treatment of human SDH tissue in either saline or BDNF. *Right:* Average KCC2 intensity values from SDH neurons of saline (light green) versus BDNF-treated (dark green) spinal segments of 12 female human donors, with comparisons of membrane and intracellular regions below. **C and D)** Representative confocal images of male (C) and female (D) human superficial dorsal horn incubated in saline or BDNF. KCC2 (red), CGRP (green) and DAPI (blue). A zoomed region (top right) shows a neuron expressing KCC2 together with the delineation of the membrane and the distance to the membrane of each pixel analysed in a colour-coded distance map. KCC2 intensity (i.u.) versus distance to the membrane profile (bottom). **E)** The *ex vivo* BDNF treatment model elicits no effect in human female SDH synaptosomes. Plots (left) and representative western blots (right) from SDH synaptosomes of human female spinal cord treated with either control saline (lilac, n=6) or 100 ng/mL recombinant BDNF for 70 minutes (purple, n=6). *p<0.05

**Figure 4.**
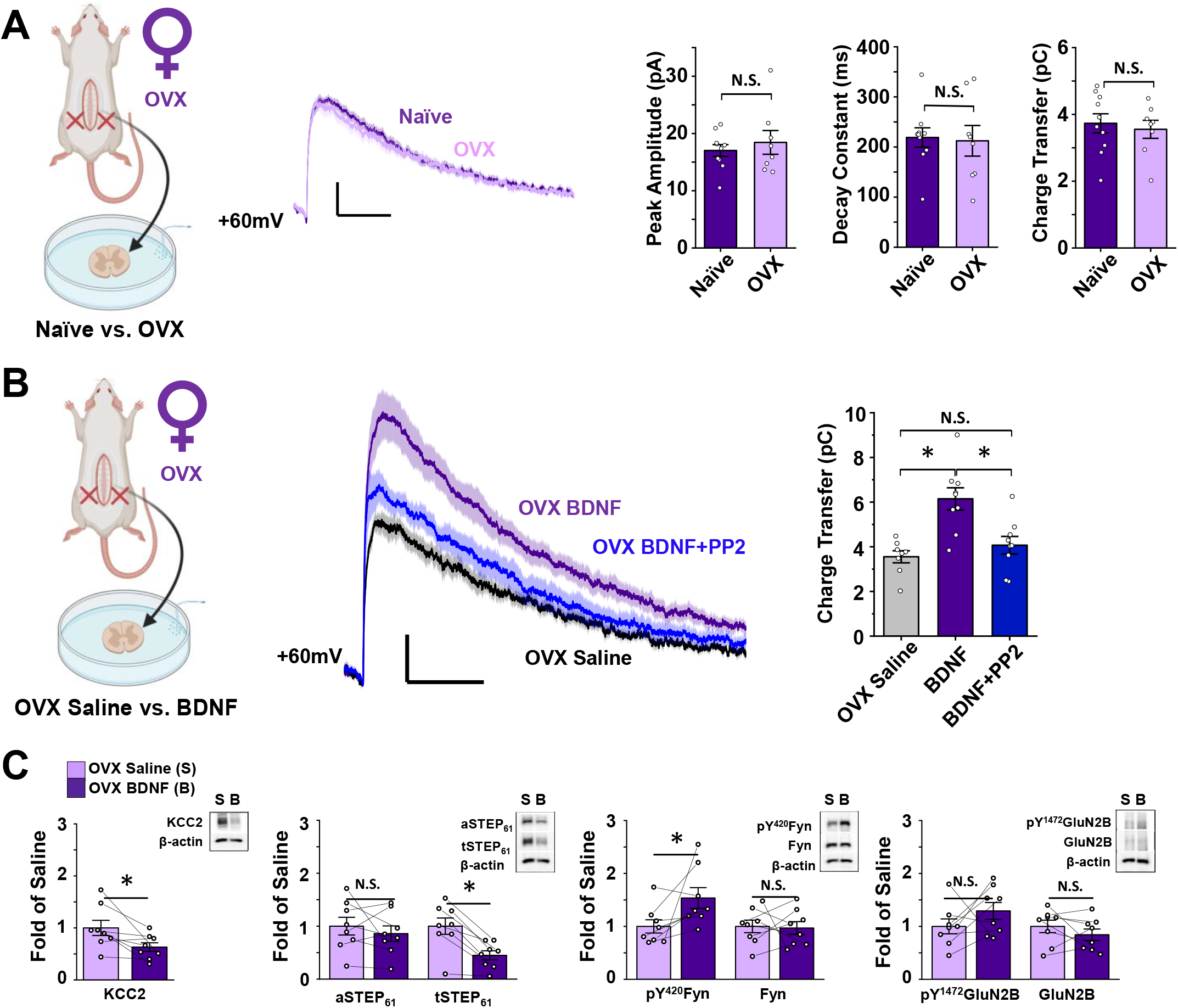
Ovariectomy triggers BDNF-mediated NMDAR potentiation by the KCC2/STEP_61_/SFK pathway in SDH neurons of female rats. **A)** Ovariectomy (OVX) has no effect on baseline lamina I NMDAR mEPSCs of female SD rats. *Left:* saline treated naïve female vs. OVX female lamina I neurons. *Middle:* Average mEPSCs at +60 mV in lamina I neurons of naïve (purple) and ovariectomized (lilac) female rats. *Right:* peak amplitude, decay constant and charge transfer of the NMDAR component of mEPSCs do not differ between naïve (n=10) and OVX (n=8) female rats. **B)** In OVX rats, NMDAR mEPSCs in lamina I neurons are potentiated following *ex vivo* BDNF treatment. This potentiation is blocked using co-treatment with the SFK inhibitor, PP2. *Left:*Recordings from lamina I neurons were compared for saline-treated versus BDNF-treated slices from OVX rats. *Middle:* Average mEPSCs at +60 mV from OVX female rat lamina I neurons; control in black, BDNF in purple, BDNF+PP2 in blue. *Right:* charge transfer of NMDAR mEPSCs shown on left. n=8 for control, n=9 for BDNF and BDNF + PP2. **C)** *Ex vivo* BDNF treatment in OVX rat SDH synaptosomes results in upregulation of pY^420^Fyn and downregulation of KCC2 and STEP_61_. Plots (left) and representative western blots (right) from OVX female rat SDH synaptosomes of tissue treated with either control saline (lilac, n=8) or 50 ng/mL recombinant BDNF for 70 minutes (purple, n=8). *p < 0.05

Comparison of means for Figure 3A-D was carried out using GraphPad Prism 9. To compare KCC2 membrane values, a region ranging from - 0.5 μm to + 0.5μm from the KCC2 peak intensity was taken. Gaussian curves were determined as the best equations to characterize this region. Amplitudes, means and standard deviations were compared using the extra sum-of-squares F-test method. For males KCC2 membrane intensities, two different Gaussian curves were required to fit the values (P =0.0002). For females KCC2 membrane intensities, two Gaussian curves were not requited to fit the values (P=0.2551).

KCC2 Membrane fit: Gaussian distribution

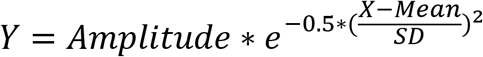

with Y = I_KCC2_; KCC2 intensity; and X is the intensity of KCC2 for each distance from the membrane. Mean = Mean KCC2 intensity (peak of the Gaussian)

To compare KCC2 intracellular values, a region ranging from 0.6 μm to 2.5μm from the KCC2 peak intensity was taken. One phase exponential decays were determined as the best equations to characterize this region. Amplitude, means and standard deviation were compared using the extra sum-of-squares F-test method. For male KCC2 intracellular intensities, two exponential decay curves were required to fit the values (P <0.0001). For female KCC2 membrane intensities, two exponential decay curves were not requited to fit the values (P=0.7235).

KCC2 intracellular fit: exponential decay

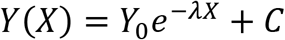

with Y = I_KCC2_ (KCC2 intensity), X = distance form the membrane, C = the baseline KCC2 intensity

For a complete list of statistical tests, comparisons made, and exact p values, see Supplementary Table 2.

## Results

### The CFA inflammatory pain model induces tactile allodynia in male and female adult rats, but only triggers changes in markers of the pathological NMDAR potentiation pathway at male SDH synapses

Despite a lack of investigation into neuronal mechanisms of pathological pain in females, there is evidence for sexual divergence in the neurophysiological features of dorsal horn pain processing. For example, we recently showed that NMDAR subunit distribution across the dorsal horn differs between sex in naïve juvenile rats^44^. We therefore investigated spinal mechanisms underlying pathological pain in male versus female adult rats using the CFA hindpaw injection model of persistent inflammatory pain^45^. Before examining markers of SDH neuronal signalling, we first tested if CFA injection induced pain hypersensitivity in both sexes by measuring behavioural pain responses. Both male and female rats that received an intraplantar injection of CFA displayed a sustained and significant reduction in PWT, while vehicle-injected rats showed no significant change in PWT (Figure 1A and C, Supplementary Table 2). The reduction in PWT persisted from one to five days post-CFA injection, demonstrating that mechanical allodynia is maintained throughout this time range in both sexes (Supplementary Figure 1).

Biochemical analysis on lumbar spinal cord tissue collected from male experimental rats revealed that CFA treatment caused a trend toward a decrease in KCC2 (p = 0.053), a marker of disinhibition, and a significant decrease in active STEP_61_ (p = 0.0085), which links disinhibition to facilitated excitation^27^, in ipsilateral SDH crude synaptosomes from CFA-injected versus vehicle-injected animals (n = 8 for each group, Figure 1B, Supplementary Figure 2, Supplementary Table 2). We also observed a significant increase in active Fyn (p = 0.013), the GluN2B-phosphorylating kinase, and significant increases in overall and phosphorylated GluN2B (p = 0.023 and p = 5.61E-4, respectively), the dominant NMDAR GluN2 subunit at adult lamina I excitatory synapses^36^, at SDH synapses of CFA-versus vehicle-injected male rats (n = 8 for all targets, Figure 1B, Supplementary Figure 2, Supplementary Table 2). These results confirm that the CFA inflammatory pain model induces the pathological KCC2/STEP_61_/Fyn-dependent GluN2B NMDAR potentiation pathway at SDH synapses of male adult rats, as we have previously reported^27^.

In sharp contrast to males, parallel experiments in females revealed no significant changes in the above markers of disinhibition (KCC2: p = 0.78; aSTEP_61_: p = 0.96; tSTEP_61_: p = 0.29) and facilitated excitation (pY^420^Fyn: p = 0.13; Fyn: p = 0.85; pY^1472^GluN2B: p = 0.60; GluN2B: p = 0.43) in the ipsilateral SDH crude synaptosome fraction from CFA-injected versus vehicle-injected female rats (n = 8 for all targets, Figure 1D, Supplementary Figure 3, Supplementary Table 2). In addition, no significant CFA-mediated changes in any of the above molecular targets were found in the crude synaptosome fractions of the remainder of the spinal cord for either sex, when compared to vehicle-treated controls (Supplementary Figures 4-7, Supplementary Table 2). The lack of changes in molecular markers of this NMDAR potentiation and SDH hyperexcitability pathway in female CFA rats suggests that neuronal mechanisms of pathological pain may diverge between male and female rats.

### Unlike males, NMDAR responses at lamina I synapses are not potentiated by *ex vivo* BDNF nor *in vivo* CFA models of pathological pain in female rats

Given the observed sex differences in molecular markers of SDH hyperexcitability, we next investigated the functional properties of excitatory synaptic NMDARs within lamina I, the outermost layer of the SDH that contains the output projection neurons of the spinal nociceptive network^46^. To test whether synaptic NMDAR responses differ between naïve male and female rats at baseline, we measured the biophysical properties of the prominent, slow NMDAR component of mEPSCs recorded from lamina I neurons held at +60 mV^36^. We found that the peak amplitude of the NMDAR component of mEPSCs, measured from 18 to 22 ms after event onset, did not significantly (p = 0.22) differ between male (I_peak_= 14.66 ± 1.54 pA, n = 10 cells from 8 animals) and female (I_peak_ = 17.03 ± 1.02 pA, n = 10 cells from 6 animals) adult rats (Figure 2A). Furthermore, we observed no sex differences in decay constant (male τ-decay = 252 ± 26 ms, female τ-decay = 220 ± 19 ms, p = 0.063), nor NMDAR charge transfer measured from 40-500 ms after event onset (male Q = 3.06 ± 0.18 pC, female Q = 3.73 ± 0.29 pC, p = 0.32) for mEPSCs recorded from lamina I neurons of male (n = 10 cells from 8 animals) versus female (n = 10 cells from 6 animals) rats (Figure 2A). These results suggest that baseline synaptic NMDAR physiology within lamina I neurons is conserved between sexes in rats.

To directly investigate whether synaptic NMDARs are differentially modulated across sex during pathological pain processing, we compared NMDAR mEPSCs in lamina I neurons from the rat *ex vivo* BDNF and *in vivo* CFA pathological pain models, in both sexes. In male rats, NMDAR charge transfer from lamina I neurons of CFA animals with validated pain hypersensitivity (PWT measured in Figure 1A) was robustly potentiated in comparison to NMDAR mEPSCs from naïve control-treated animals (p = 0.032, n = 10 cells from 8 animals for control, 8 cells from 6 animals for CFA, Figure 2B, Supplementary Table 2). Furthermore, using a previously validated rodent *ex vivo* BDNF model of pathological pain where slices from naïve rats are incubated in 50 ng/mL recombinant BDNF for a minimum of 70 minutes^25,27^, we found a significant potentiation of NMDAR responses at lamina I synapses from BDNF-versus control-treated spinal cord slices (p = 2.90E-6, n = 10 control cells from 8 animals, n = 6 BDNF cells from 5 animals, Figure 2B, Supplementary Table 2, control and CFA data originally published by Dedek *et al*^27^).

Under the same conditions and in parallel (when possible) to the experiments performed in males, we investigated the properties of lamina I synaptic NMDAR responses from control versus pathological pain model female rats. Strikingly, we found no potentiation of NMDAR responses at female lamina I synapses by either the *ex vivo* BDNF or *in vivo* CFA models of pathological pain (Figure 2C). The charge transfer through NMDAR mEPSCs from lamina I neurons of CFA-treated females was not different from NMDAR mEPSCs of control-treated animals (p = 0.25, n = 10 control cells from 6 animals, n = 8 CFA cells from 5 animals, Figure 2C, Supplementary Table 2). As was the case for males (Figure 1A), female CFA-treated animals showed significantly decreased PWT (Figure 1C), eliminating that the possibility that the lack of NMDAR potentiation was due to an absence of pain hypersensitivity in this cohort of female CFA animals. Furthermore, synaptic NMDAR responses from female spinal cord slices treated with BDNF were not significantly different from corresponding control-treated slices (p = 0.25, n = 10 control cells from 6 animals, n = 8 BDNF cells from 5 animals, Figure 2C, Supplementary Table 2), demonstrating a lack of potentiation of synaptic NMDARs by *ex vivo* BDNF treatment in female rats. A previous study has found a slower onset of BDNF-mediated behavioural hypersensitivity in female mice compared to male mice following *in vivo* intrathecal administration of BDNF^17^. To investigate whether prolonged exposure to BDNF *ex vivo* would induce NMDAR potentiation at lamina I synapses of female rats, we performed a subset of experiments using slices incubated in 50 ng/mL BDNF for 2 to 4.5 hours. As was the case in slices incubated for a minimum of 70 minutes, NMDAR charge transfer was not different between control- and prolonged BDNF-treated slices, and thus were grouped into the overall BDNF-treated female cells (p = 0.89, n = 10 control cells from 6 animals, n = 6 long-BDNF cells from 3 animals, Supplementary Figure 8, Supplementary Table 2). Furthermore, in another subset of neurons, slices were incubated in 100 ng/mL BDNF for 2+ hours and showed no difference in NMDAR charge transfer compared to control-treated recordings (data not shown). Taken together, our findings indicate that although male and female rats exhibit conserved baseline NMDAR responses, BDNF-mediated pathological pain processes drive a selective potentiation of NMDAR responses at lamina I synapses in male but not female adult rats.

We next sought to explore whether *ex vivo* BDNF treatment would alter any molecular markers of the disinhibition–STEP_61_ downregulation–NMDAR potentiation spinal hyperexcitability pathway in female rats. As was seen in the *in vivo* CFA inflammatory pain model for females (Figure 1D), *ex vivo* treatment of SDH tissue with recombinant BDNF did not elicit significant changes in a marker of disinhibition, KCC2 (p = 0.88), in active or total STEP_61_ (p = 0.74, p = 0.47, respectively) or markers of increased excitability (pFyn p = 0.64, Fyn p = 0.60, pGluN2B p = 0.70, GluN2B p = 0.62) in female rat crude synaptosomes compared to paired SDH treatment with saline control (Figure 2D, Supplementary Figure 9, Supplementary Table 2, n = 8 for all targets). This is in stark contrast to male rats, where we previously found that *ex vivo* BDNF treatment mediated a significant downregulation of KCC2, active STEP_61_ and total STEP_61_, and an upregulation of pFyn, pGluN2B and GluN2B^27^. As observed in both CFA-treated male tissue (Figure 1B) and previously in the male *ex vivo* BDNF pathological pain model^27^, we also found no CFA/BDNF-mediated changes in the above molecular markers in the synaptosome fractions of the remainder of the spinal cord (Supplementary Figures 10-11, Supplementary Table 2) of female rats. From these data, we conclude that, unlike in males, *ex vivo* BDNF treatment does not induce the pathological KCC2-STEP_61_-Fyn-GluN2B signalling pathway at SDH synapses of female rats.

### *Ex vivo* BDNF treatment results in sexually dimorphic effects on markers of disinhibition and facilitated excitation at SDH synapses of human spinal cord tissue

A major barrier to the development of novel therapeutic approaches for pain is a lack of human preclinical models^32,47^. To address this, we have previously developed a human *ex vivo* BDNF model of pathological pain using tissue from human organ donors^27^. In this model, isolated spinal cord tissue was collected from adult neurological determination of death organ donors and incubated in oxygenated solution containing either 100 ng/mL recombinant BDNF or a saline control for 70 minutes, similar to that done for rats^27^. Here, we first probed for a marker of BDNF-mediated disinhibition in SDH neurons using KCC2 immunostaining at membrane versus intracellular compartments^27,48^, with experiments on spinal cord tissue from male and female donors done in parallel. CGRP immunostaining was used to identify the SDH^49^. Consistent with our initial findings^27^, experiments with an expanded male sample size demonstrated that *ex vivo* BDNF treatment mediates a significant decrease in membrane KCC2 intensity (p = 2.09E-4) and a significant increase in intracellular KCC2 intensity (p = 1.75E-4) in CGRP-positive SDH neurons of male human donors when compared to adjacent saline-treated control tissue (n = 12, Figure 3A,C, Supplementary Table 2). In female humans, however, we saw no change in KCC2 intensity at neuronal membranes or intracellular compartments following BDNF treatment (p = 0.26, p = 0.72, n = 10, Figure 3B,D, Supplementary Table 2), demonstrating that *ex vivo* BDNF treatment does not drive an internalization of KCC2 from female SDH neuronal membranes. This suggests that unlike in males, *ex vivo* treatment with BDNF does not result in KCC2-dependent disinhibition in SDH neurons of human females.

To further test whether *ex vivo* BDNF activates the pathological disinhibition to facilitated excitation pathway in male versus female human SDH neurons, we next probed synaptic proteins using western blot analysis. In SDH crude synaptosomes of human males, we have previously shown that BDNF treatment triggers downregulation of KCC2 and total and active STEP_61_, indicating disinhibition, as well as upregulation of phosphorylated Fyn and phosphorylated GluN2B, markers of facilitated excitation^27^. With sufficient samples now collected from female human donors, which were less frequent but collected and processed in parallel with male donor tissue, here, we tested the effects of *ex vivo* BDNF on the same molecular markers in female SDH neurons. Compared to saline-treated controls, treatment of female human spinal cord tissue with BDNF had no effect on markers of disinhibition (KCC2 p = 0.47, aSTEP_61_ p = 0.60, tSTEP_61_ p = 0.77) nor on markers of facilitated excitation (pFyn p = 0.49, Fyn p = 0.63, pGluN2B p = 0.60, GluN2B p = 0.82) in the SDH crude synaptosome fraction (n = 6, Figure 3E, Supplementary Figure 12, Supplementary Table 2). Similar to male and female rats, no changes in these targets were seen in the crude synaptosome fraction from the remainder of the spinal cord as a result of BDNF treatment, except for a change in aSTEP_61_ (p = 0.025, n = 6, Supplementary Figures 13-14, Supplementary Table 2). The above complementary KCC2 internalization and synaptic marker assays provide compelling evidence that BDNF selectively drives a coupling between KCC2-dependent disinhibition, STEP_61_ downregulation and GluN2B NMDAR potentiation in human SDH neurons of males but not females. We therefore conclude that the sexual dimorphism in this spinal molecular mechanism underlying pathological pain is conserved between rats and humans.

### Ovariectomy recapitulates the BDNF – disinhibition –NMDAR potentiation pathway in SDH neurons of female rats

In a final series of experiments, we aimed to understand the biological underpinnings for the lack of the BDNF-driven coupling between disinhibition and facilitated excitation in female SDH neurons. Since sexually dimorphic NMDAR signalling in pain modulation has previously been found to be controlled by female sex hormones^50^, we explored whether removal of female sex hormones during development would alter sensitivity to BDNF at adult female SDH synapses. We used female rats that were ovariectomized on postnatal day 21, before reaching sexual maturity, and were then left to develop to the adult ages used throughout our animal studies (3 to 4 months old). We first compared the effects of ovariectomy (OVX) on baseline lamina I synaptic NMDAR properties in adult rats. As observed for the comparisons of lamina I NMDAR mEPSCs at +60 mV between male and female rats (Figure 2A), we found no significant (p = 0.97) difference in NMDAR mEPSC peak amplitude between age-matched naïve (I_peak_ = 17.0 ± 1.0 pA, n = 10 cells from 6 animals) and OVX (I_peak_ = 18.4 ± 2.1 pA, n = 8 cells from 5 animals) adult female rats (Figure 4A). Similarly, OVX treatment did not alter the charge transfer (naïve Q = 3.73 ± 0.29 pC, OVX Q = 3.55 ± 0.27 pC, p = 0.66) and decay constant (naïve τ-decay = 220 ± 19 ms, OVX τ-decay = 212 ± 31 ms, p = 0.85) of the NMDAR component of mEPSCs in lamina I neurons from age-matched female rats (Figure 4A, Supplementary Table 2). These findings suggest that removal of female sex hormones during early development does not significantly change the overall NMDAR subunit-driven^36^ biophysical properties of NMDAR responses at adult lamina I synapses under baseline conditions.

Next, we tested the effects of *ex* vivo BDNF treatment on synaptic NMDAR responses in lamina I neurons from OVX adult rats. Surprisingly, we found that NMDAR mEPSCs were robustly potentiated by BDNF treatment in OVX animals, with a significant (p = 5.01E-4) increase in NMDAR charge transfer for OVX rat slices incubated in BDNF (n = 9 cells from 7 animals) compared to OVX slices incubated in saline control solution (n = 8 control cells from 5 animals; Figure 4B, Supplementary Table 2). Previously, we have shown that PP2, a SFK inhibitor^51^, blocks BDNF-mediated potentiation of GluN2B-containing NMDARs at male rat lamina I synapses^25^. Here, we tested whether co-treatment with PP2 would attenuate the BDNF-mediated potentiation of synaptic NMDAR responses in spinal slices from OVX animals. As hypothesized, synaptic NMDAR charge transfer for OVX lamina I neurons treated with both BDNF and 1 μM PP2 did not differ from OVX control cells (p = 0.65, n = 8 OVX control cells from 5 animals, n = 9 BDNF+PP2 cells from 5 animals), but instead significantly differed from NMDAR responses of OVX lamina I neurons treated with BDNF alone (p = 0.0034, n = 9 cells from 7 animals, Figure 4B, Supplementary Table 2). Thus, unlike in naïve female rats (Figure 2C) but similar to male rats^25^, *ex vivo* BDNF treatment drives an SFK-dependent potentiation of NMDAR responses at lamina I synapses in female rats that have their sex hormones removed early in development.

Lastly, we tested the effects of OVX treatment on the potential BDNF-mediated disinhibition to facilitated excitation molecular mechanism at SDH synapses. In contrast to that found for naïve female rats (Figure 2D) and female humans (Figure 3E), we found that *ex vivo* BDNF treatment induced significant changes in several proteins linked to the disinhibition/facilitated excitation pathway in SDH synaptosomes from OVX rats (Figure 4C): BDNF treatment resulted in downregulation of KCC2 (p = 0.040) and total STEP_61_ (p = 6.16E-4), and upregulation of pY^420^Fyn (p = 0.046) (no changes in: aSTEP_61_: p = 0.41, Fyn: p = 0.87, pY^1472^GluN2B: p = 0.24, and GluN2B: p = 0.33; n = 8 for all targets, Figure 4C, Supplementary Figure 15, Supplementary Table 2). In the crude synaptosome fractions from the remainder of the spinal cord, we observed significant changes in Fyn (p = 0.011) and total STEP_61_ (p = 0.0014; Supplementary Figures 16-17, Supplementary Table 2), suggesting that OVX alters the response to BDNF not only at SDH synapses, but throughout the entire spinal cord. Together, these findings suggest that female rats ovariectomized before reaching sexual maturity respond in a male-like manner to BDNF treatment, switching from the non-BDNF-sensitive naïve female phenotype to the male disinhibition-to-NMDAR-potentiation phenotype in SDH neurons.

## Discussion

Given the importance of understanding the neurobiological underpinnings of chronic pain across sex, we investigated mechanisms of neuronal hyperexcitability in the rat and human superficial dorsal horn of males and females. Using the *in vivo* CFA hindpaw injection model of inflammatory pain, we found that although both sexes of rat displayed tactile allodynia, the STEP_61_-pFyn-pGluN2B spinal hyperexcitability pathway was activated in males, but not in females. Unlike males, lamina I synaptic NMDAR responses were not potentiated in CFA-injected female rats. Moreover, *ex vivo* BDNF treatment of spinal cord slices did not potentiate NMDAR mEPSCs nor drive the pathological coupling mechanism between disinhibition and facilitated excitation at SDH synapses of female rats, as is observed for males. Building on our previous work establishing the *ex vivo* BDNF model of pathological pain in male humans^27^, we tested the effects of *ex vivo* BDNF treatment on female human SDH tissue. We found that, like female rats, female human SDH tissue showed no BDNF-driven changes in the distribution and expression of synaptic proteins linked to hyperexcitability, with no internalization of KCC2 from neuronal membranes. Taken together, our female rat and human experiments show that BDNF does not drive the pathological KCC2/STEP_61_/Fyn-mediated potentiation of synaptic NMDARs within the female SDH, as it does in males. We conclude that this sex difference in response to BDNF is hormonally-mediated, with female rats ovariectomized before reaching sexual maturity displaying a male-like response to BDNF characterized by SFK-dependent NMDAR potentiation and associated changes in synaptic markers of disinhibition and facilitated excitation.

Although sex differences in immune modulation of pain have been recently discovered^5,14^, it has been more challenging to study potential sexual dimorphism in neuronal nociceptive signalling. Based on pharmacological evidence where blocking neuronal targets *in vivo* alleviates behavioural hypersensitivity in both sexes, it has been assumed that pathological pain mechanisms are conserved once they reach the neuronal level. For example, the NMDAR antagonist APV blocks nerve injury-induced allodynia in both sexes^13^ and so it has been proposed that NMDAR-mediated facilitated excitation is conserved from males to females^11^. However, this analgesic efficacy of NMDAR antagonists only demonstrates that NMDARs are critical determinants of excitability in pain processing neurons for both sexes^52^, and not that NMDARs themselves are dysregulated to specifically mediate pathological pain across sex. This critical distinction is easy to overlook, given the overwhelming evidence from decades of research using male or unsexed animals implicating NMDARs as molecular targets for spinal hyperexcitability and inflammatory pain^27,53–55^. Although we found that baseline synaptic NMDAR responses are conserved between sex, we observed a dramatic sexual divergence in NMDAR dysregulation during pathological pain processing, with potentiation of lamina I synaptic NMDAR responses in inflammatory and *ex vivo* BDNF pain models for males but not females. Sorge, Mapplebeck and colleagues have shown that the development of nerve injury-induced mechanical allodynia is impaired in BDNF-null males, but not females, suggesting that BDNF is not required for the development of tactile allodynia in female mice^13^. However, it remains unknown if NMDARs are potentiated in the SDH in female rodent models of neuropathic pain, and future studies are needed to address this important question.

In contrast to the lack of evidence for pathological potentiation of synaptic NMDAR responses in females, the specificity of BDNF-mediated disinhibition to only males appears to vary by the specific type of pain and underlying mechanism involved. Previous *in vivo* pharmacological studies have shown that intrathecal administration of recombinant BDNF in female rodents can induce behavioural hypersensitivity in mice and rats^17,56^ including an associated downregulation of dorsal horn KCC2^56^. In particular, Mapplebeck and colleagues showed that intrathecal BDNF administration results in tactile allodynia in both sexes, but with a longer time required for the sensitizing effect in females^17^. For this reason, we treated a subset of female rat spinal slices with BDNF for 2-4.5 hours (Supplementary Figure 8) and found no difference in NMDAR mEPSC charge transfer between the long-BDNF incubation or control-treated mEPSCs. We, therefore, propose that BDNF induces other signalling pathways such as trophic^57,58^ or immune mechanisms *in vivo* that rely on factors outside our SDH mechanism under study. It has also been found that neuropathic pain can trigger KCC2-dependent disinhibition in both sexes, including in nerve injury^17^ and diabetic neuropathy^56^ models of neuropathic pain in female rodents. Given that many neurochemical factors mediating allodynia differ between neuropathic and inflammatory pain^53,59,60^ it will be critical to investigate which molecular elements of the KCC2-STEP_61_-pFyn-pGluN2B SDH hyperexcitability pathway are altered in neuropathic pain states in females.

More broadly, our findings highlight the paucity of work relating to the molecular drivers of spinal pain pathology in females. Spinal hyperexcitability mediates pathological pain in females. Both female SNI rats^61^ and human fibromyalgia patients^62^ exhibit a decrease in nociceptive flexion reflex thresholds, which is an indirect readout of central sensitization^63^. Windup, as measured by electromyography^64^ and *in* vivo electrophysiology^65^, has also been observed in female animals with spinal cord injury. Despite this evidence, the molecular mechanisms underlying SDH hyperexcitability in females remain unclear. One possible candidate is CGRP, which has been shown to alter excitatory postsynaptic responses within higher-order pain processing circuits of the amygdala^66^ as well as in SDH neurons^67^. The roles of CGRP in mediating pain pathology may also be sexually dimorphic. In low doses, dural CGRP induces pain hypersensitivity only in females in rodent preclinical models of migraine^68^. Recent developments show that this female-specific effect of CGRP may be dependant on prolactin^69^ and involves KCC2-dependent disinhibition^70^. Further studies are needed to understand the role of CGRP in mediating the hyperexcitability of SDH neurons in males and females.

Our finding that ovariectomy results in a switch from the female BDNF-insensitive phenotype to a male-like BDNF-sensitive phenotype in SDH neurons suggests that female sex hormones have an organizational effect on spinal mechanisms of pain signalling, with persistent effects lasting into adulthood. Further study is needed to identify which female sex hormones regulate this dimorphism by isolating the effects that estrogen^71–73^ and progesterone^74^ have on the development of the nociceptive pathway in both rodents and humans. From a translational perspective, further human spinal tissue studies are also needed to determine what effects the above pathological signalling pathways have on both subnetwork functions and the overall output of the human SDH pain processing matrix. With exciting recent advances in voltage-sensitive dye imaging^75^, multielectrode array recording, and single-cell sequencing^76^ approaches, researchers can now investigate the impact of dynamic and degenerate signalling pathways on human spinal pain processing^32,77^, including well-designed parallel experiments in both males and females. Our results therefore highlight the critical need for foundational studies using both sexes^6,7^, as well as the importance of using human preclinical models to improve translation between basic science and clinical medicine in the field of pain research and beyond.

## Acknowledgements

We thank the organ and tissue donors and their families for their generous, selfless gift. We thank the Trillium Gift of Life Network, the surgical staff at The Ottawa Hospital Civic Campus, and Dr. Suzan Chen, Lei Zhou, Santina Temi, Dr. Christopher Rudyk, Ahmad Galuta, Christopher Dedek, and Dr. Sara Ameri for their help with human spinal cord collection.

## Funding

This study was supported by a John R. Evans Leaders Fund grant from the Canada Foundation for Innovation (M.E.H.), a Discovery Grant from the Natural Sciences and Engineering Research Council of Canada (M.E.H.), an Early Career Research Grant from the International Association for the Study of Pain (M.E.H.), an Early Career Investigator Pain Research Grant from the Canadian Pain Society and Pfizer Canada (M.E.H.), and Project/Operating Grants from the Canadian Institutes of Health Research (M.E.H., 388432; Y.D.K., FDN-159906).

## Notes

### Competing Interest Statement

The authors have declared no competing interest.

