## Supplementary for "Sexual dimorphism in a neuronal mechanism of spinal hyperexcitability across rodent and human models of pathological pain"

### Supplementary Figures and Tables

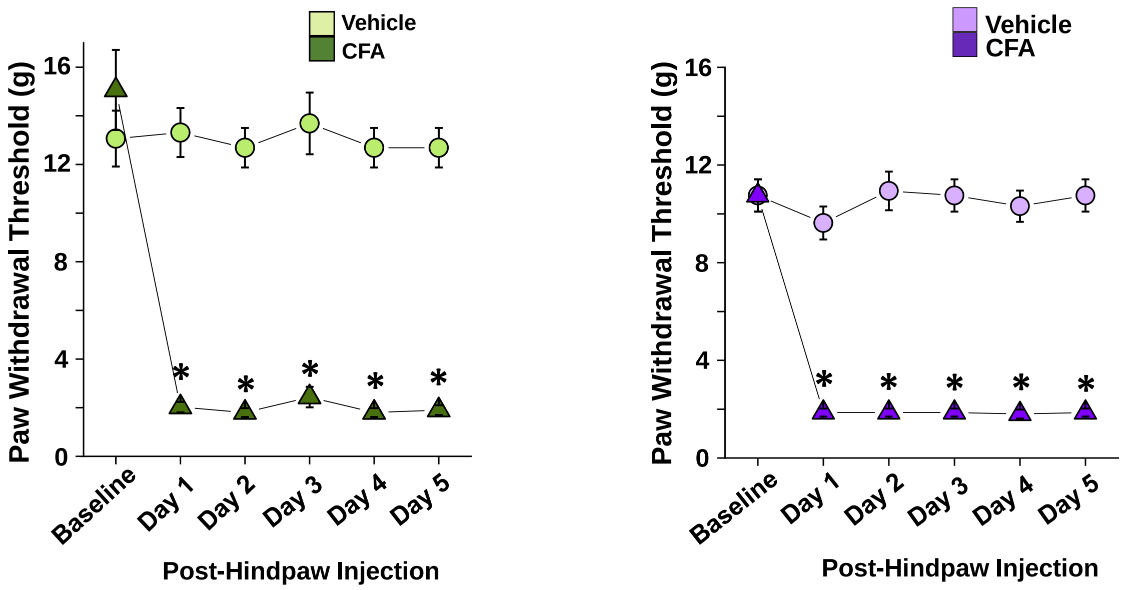


**Supplementary Figure 1. Paw withdrawal threshold is decreased in male and female rats receiving CFA.** Male rats, left, and female rats, right, that receive CFA injection display significantly decreased paw withdrawal threshold 1 – 5 days post-injection. Tissue from animals in this figure was used in Figure 1B and 1D. N = 8 animals per sex per group *$p < 0.05$.

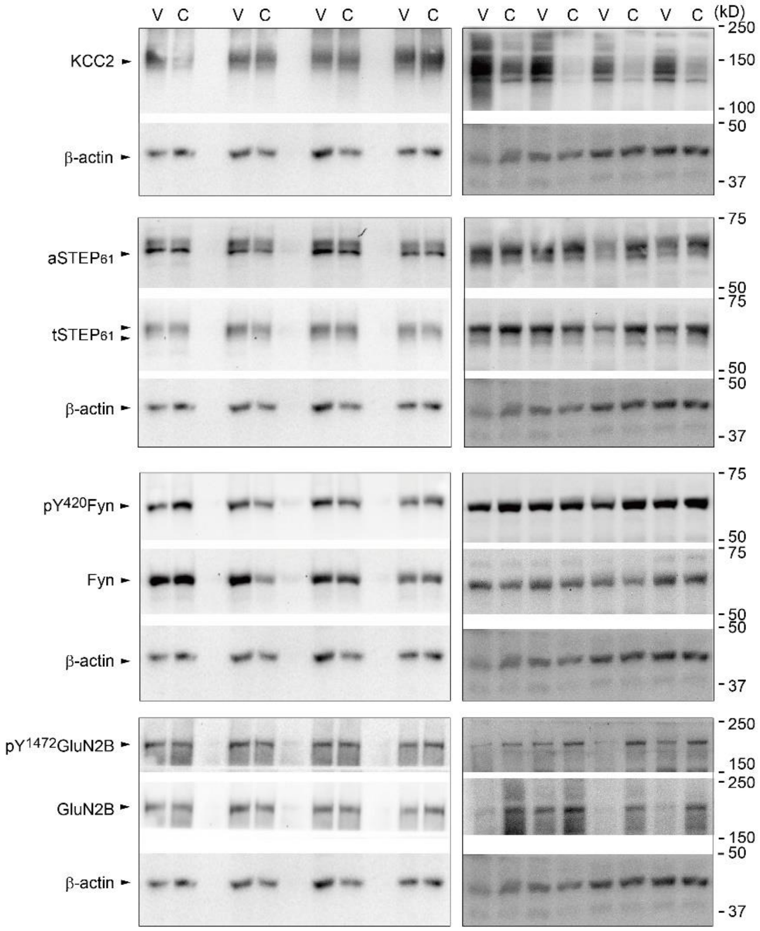


**Supplementary Figure 2. Male CFA SDH synaptosome gels from animals treated with either vehicle (V) or CFA (C).** Individual gels were cut into sections to allow for probing several targets concurrently β-actin, the loading control, can be seen under each set of targets (labelled on the left while weight of the

target, in kD is on the right. n= 8

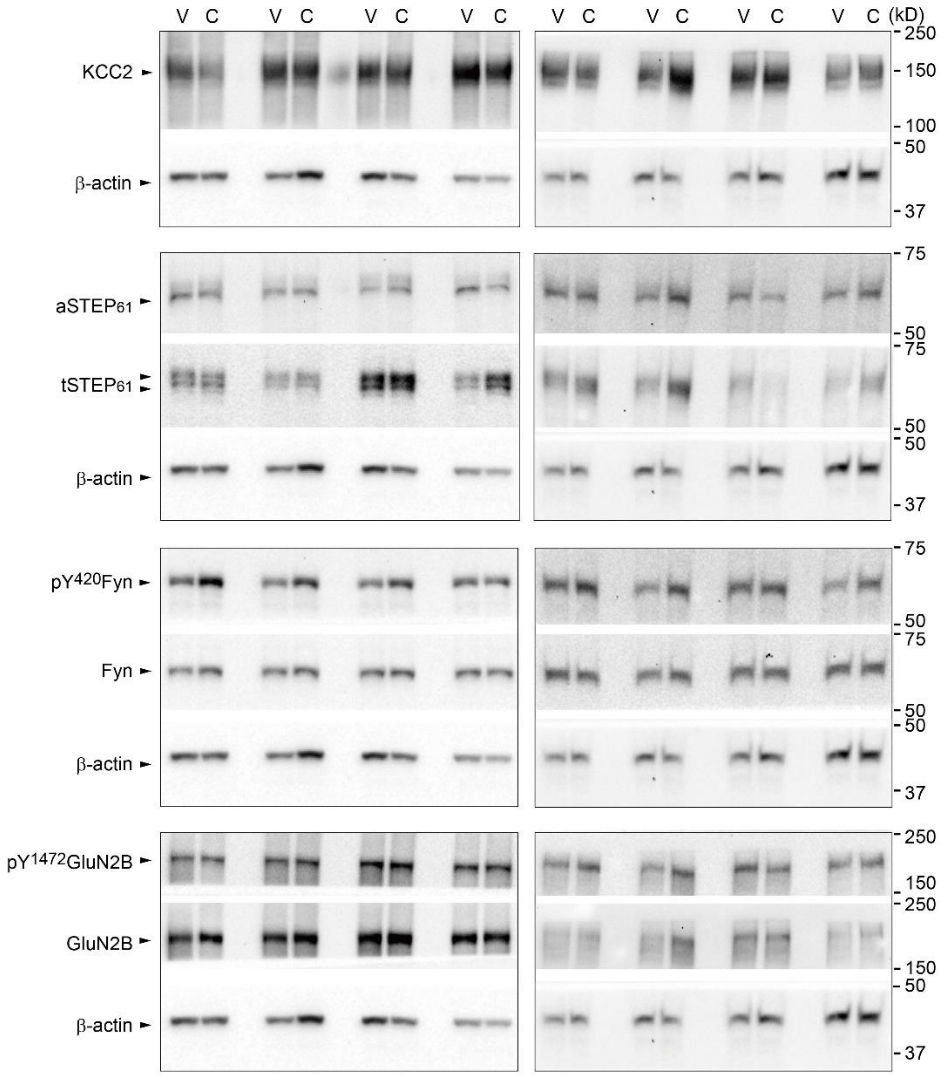


**Supplementary Figure 3. Female CFA SDH synaptosome gels from animals treated with either vehicle (V) or CFA (C).** Individual gels were cut into sections to allow for probing several targets concurrently β-actin, the loading control, can be seen under each set of targets (labelled on the left while weight of the target, in kD is on the right. n= 8

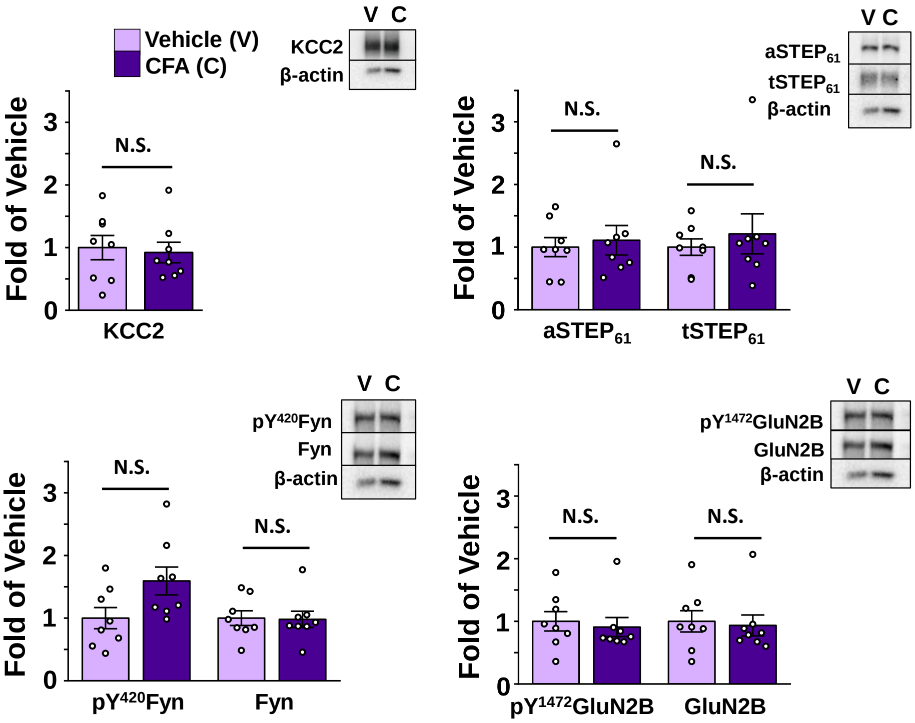


**Supplementary Figure 4. The CFA model of inflammatory pain elicits no change in our targets in crude synaptosome fractions of the portion of the spinal cord just ventral to the SDH in female rats.** Vehicle in lilac, CFA in dark purple; n = 8 animals per group.

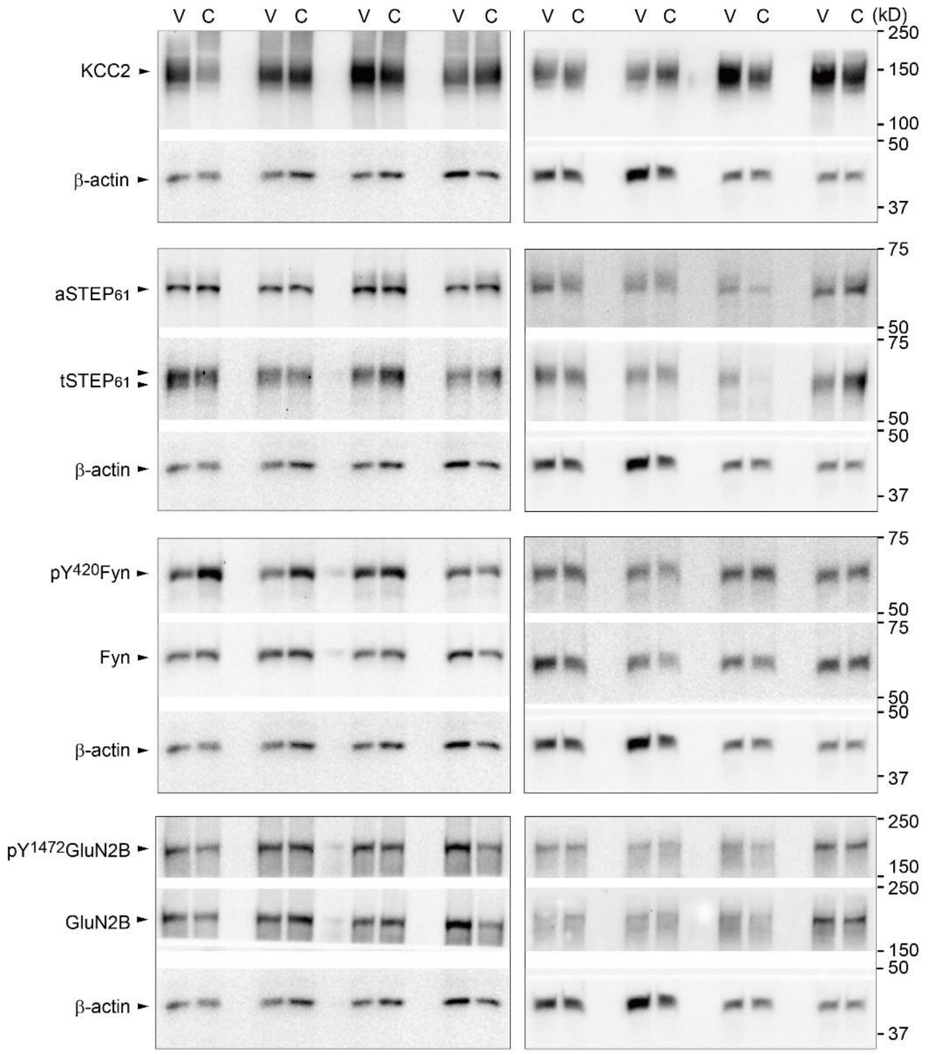


**Supplementary Figure 5. Female CFA VH synaptosome gels from animals treated with either vehicle (V) or CFA (C).** Individual gels were cut into sections to allow for probing several targets concurrently β-actin, the loading control, can be seen under each set of targets (labelled on the left while weight of the target, in kD is on the right. n= 8.

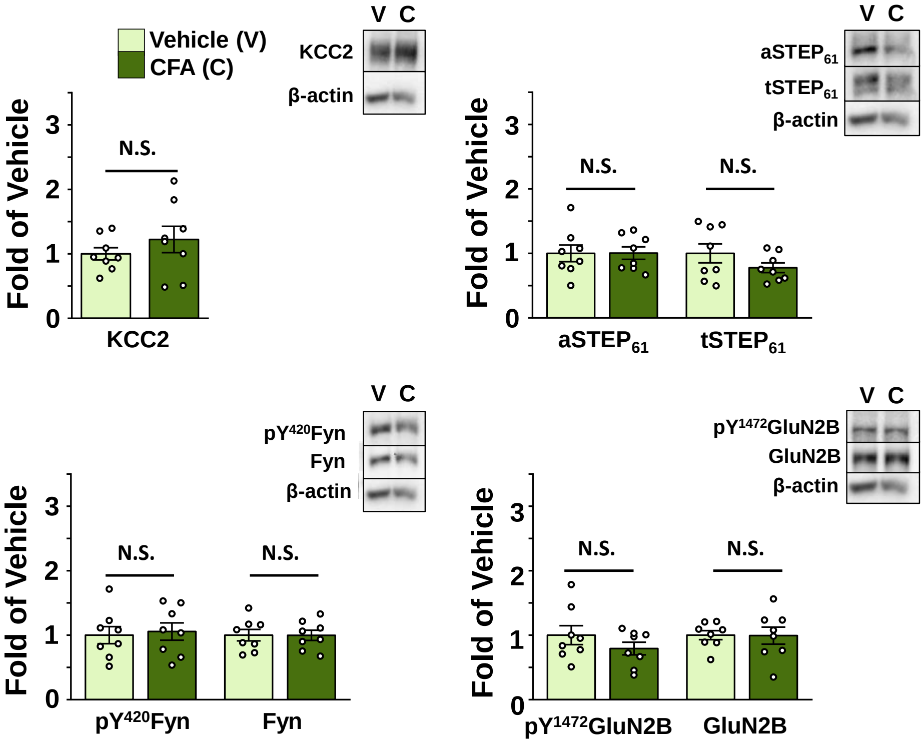


**Supplementary Figure 6. The CFA model of inflammatory pain elicits no change in our targets in crude synaptosome fractions of the portion of the spinal cord just ventral to the SDH in male rats.** Vehicle in light green, CFA in dark green; n = 8 animals per group.

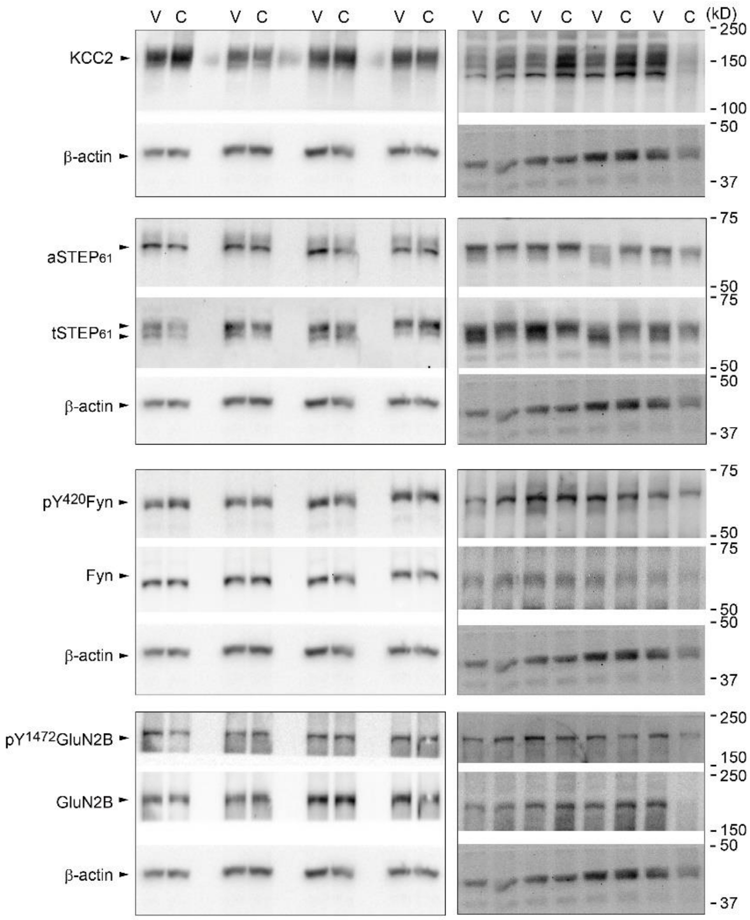


**Supplementary Figure 7. Male CFA VH synaptosome gels from animals treated with either vehicle (V) or CFA (C).** Individual gels were cut into sections to allow for probing several targets concurrently β-actin, the loading control, can be seen under each set of targets (labelled on the left while weight of the target, in kD is on the right. n= 8

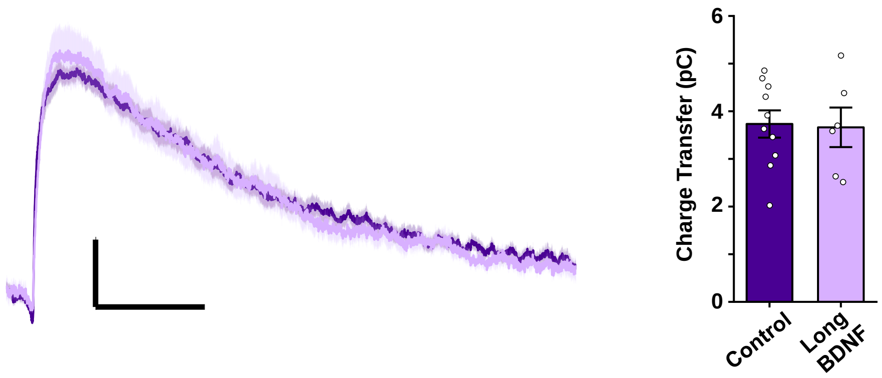


**Supplementary Figure 8. BDNF incubation for 2-4 hours does not result in potentiation of female rat lamina I mEPSCs.** Lamina I mEPSCs from female rat tissue incubated for 2-4 hours in 50ng/mL BDNF (lilac) show no change in charge transfer when compared to control-treated slices (dark purple). N =10 cells from 5 animals for control, 6 cells from 3 animals for long-BDNF.

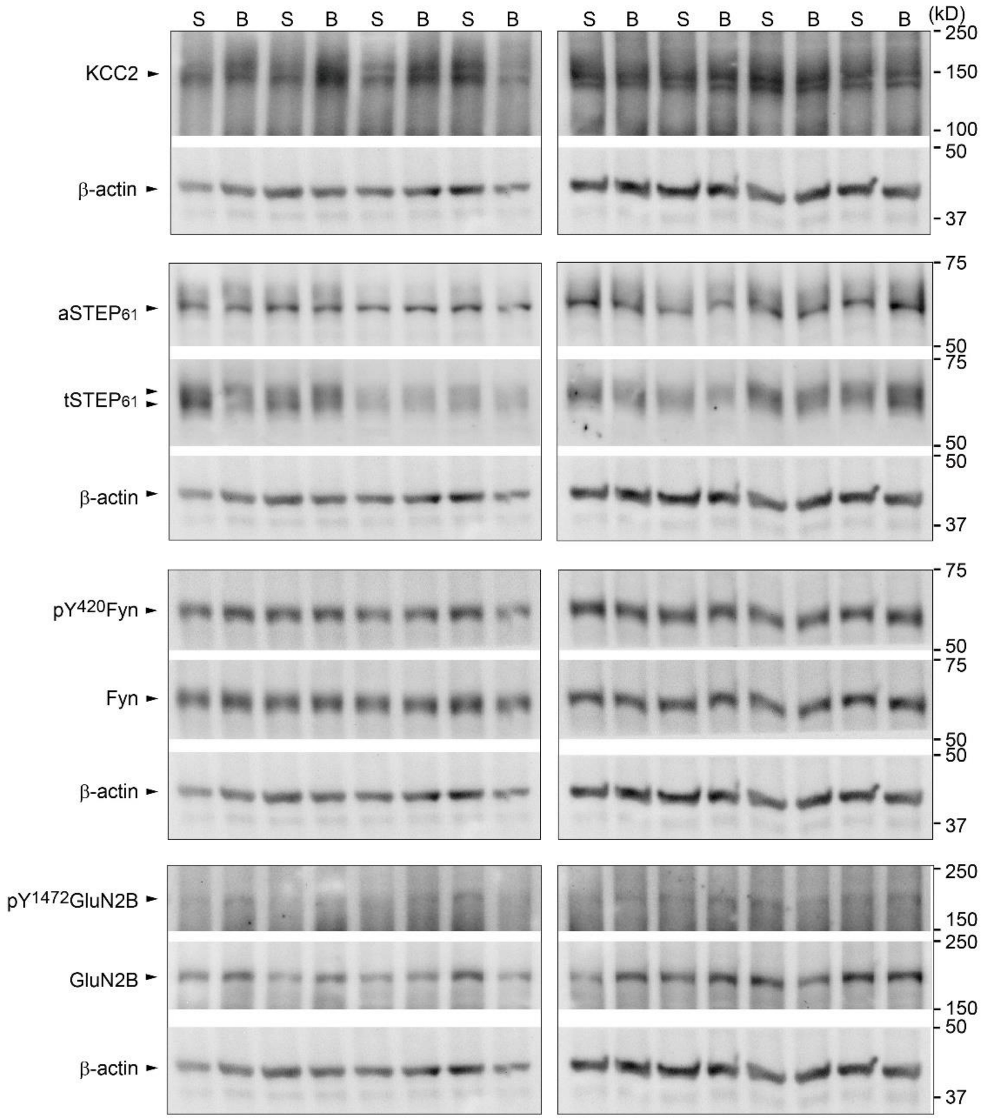


**Supplementary Figure 9. Female BDNF SDH synaptosome gels from animals treated with either saline (S) or BDNF (B).** Individual gels were cut into sections to allow for probing several targets concurrently β-actin, the loading control, can be seen under each set of targets (labelled on the left while weight of the target, in kD is on the right. n= 8

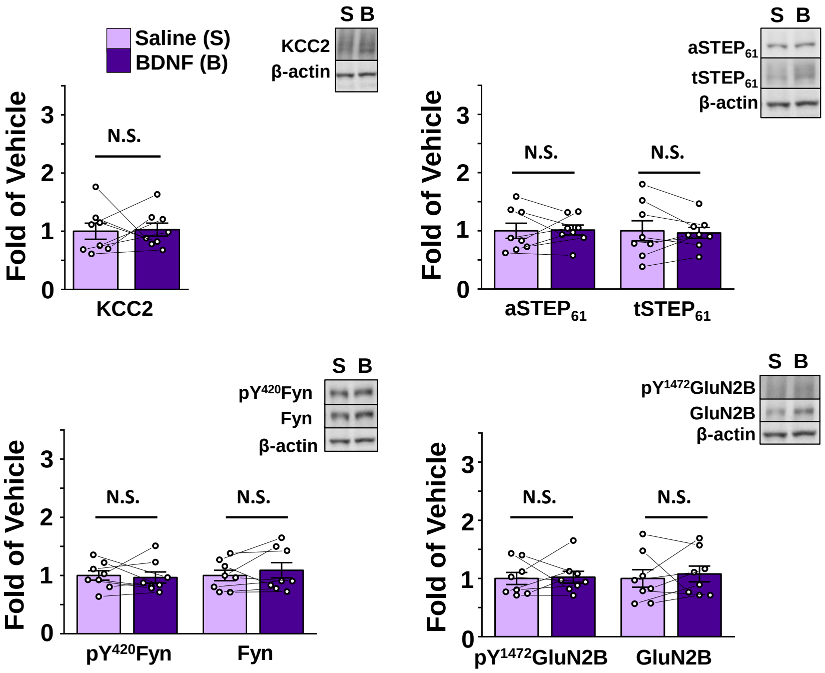


**Supplementary Figure 10. Incubating female rat tissue in 50 ng/mL BDNF elicits no change in our targets in crude synaptosome fractions of the portion of the spinal cord just ventral of the SDH**. Saline in lilac, BDNF in dark purple; n = 8 animals per group.

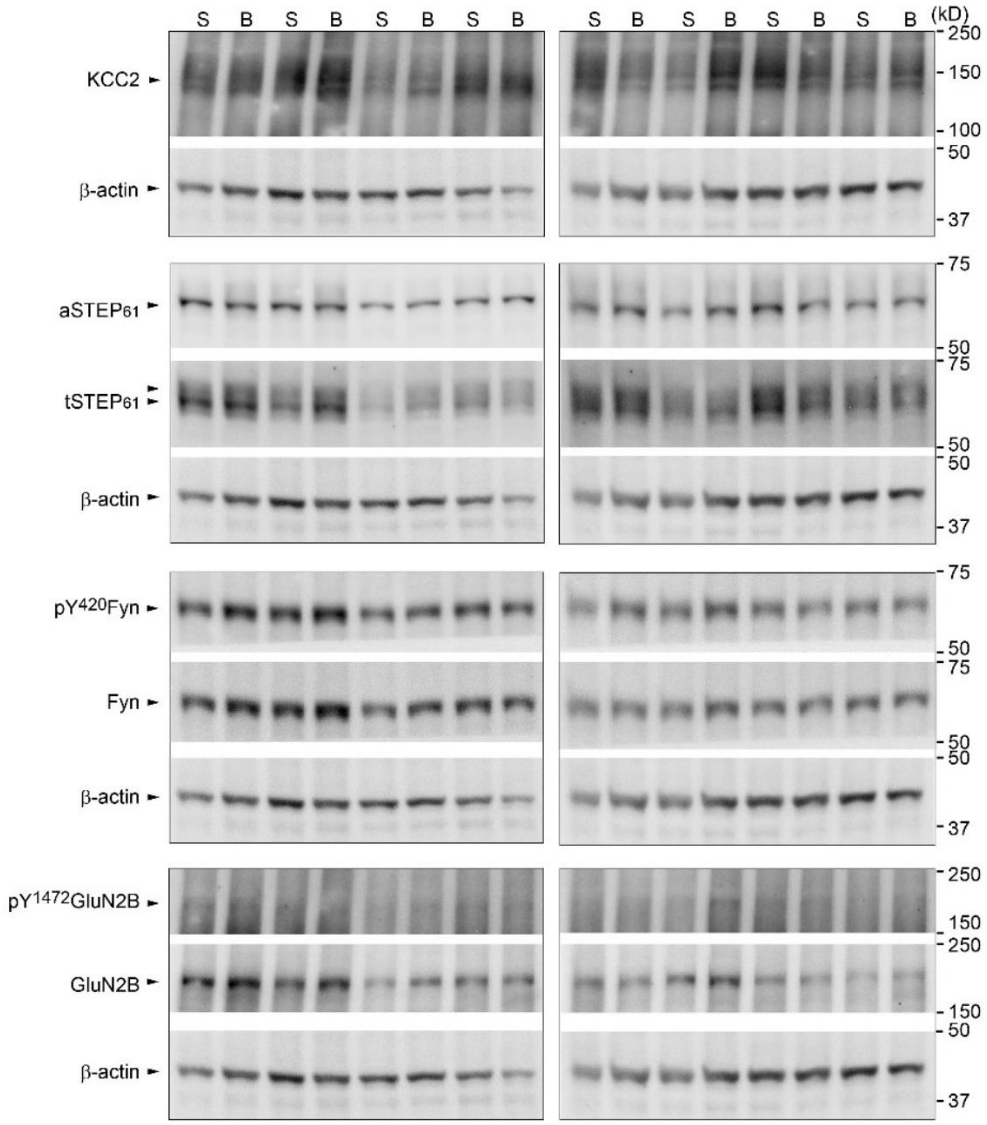


**Supplementary Figure 11. Female BDNF VH synaptosome gels from animals treated with either saline (S) or BDNF (B).** Individual gels were cut into sections to allow for probing several targets concurrently β-actin, the loading control, can be seen under each set of targets (labelled on the left while weight of the target, in kD is on the right. n= 8

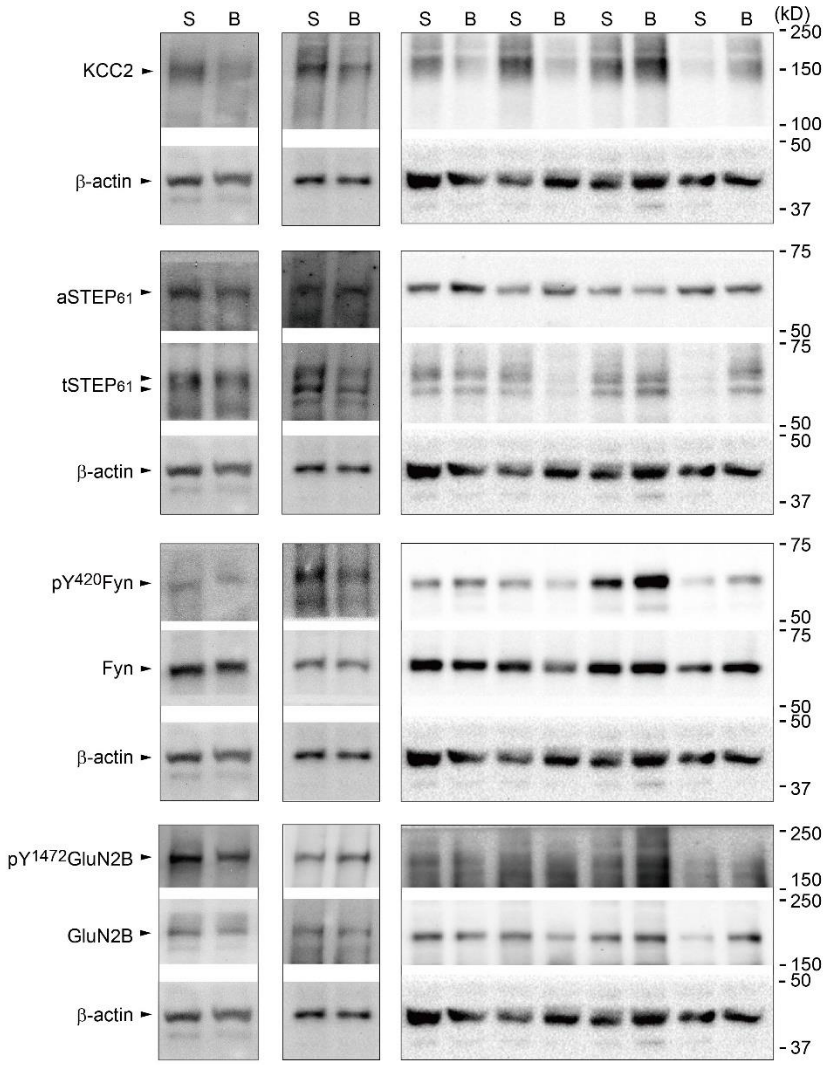


**Supplementary Figure 12. Female BDNF human SDH synaptosome gels from human spinal samples treated with either saline (S) or BDNF (B).** Individual gels were cut into sections to allow for probing several targets concurrently β-actin, the loading control, can be seen under each set of targets (labelled on the left while weight of the target, in kD is on the right. n= 6

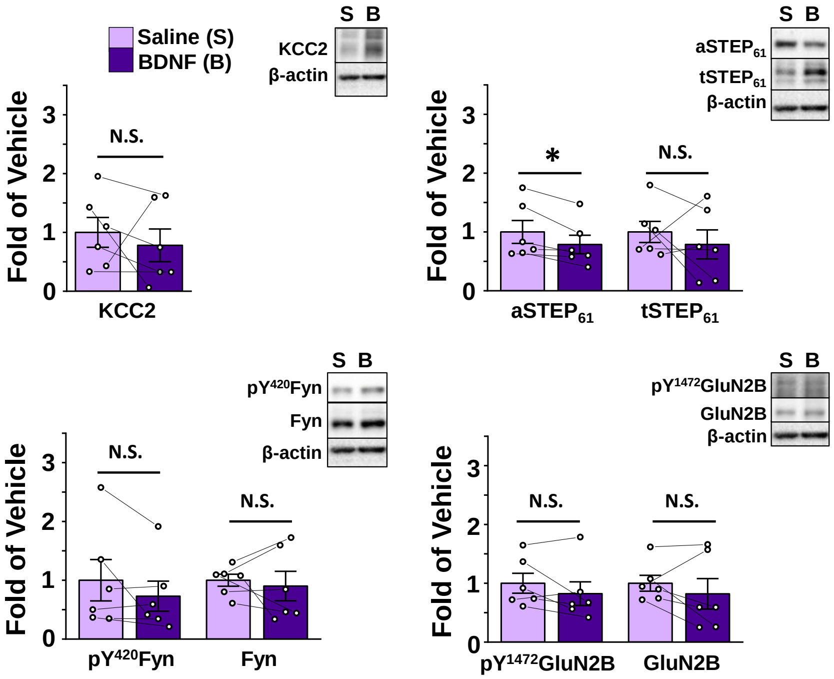


**Supplementary Figure 13. Incubating female human tissue in 100 ng/mL BDNF elicits no change in KCC2, tSTEP, pFyn, Fyn, pGluN2B, or GluN2B in crude synaptosome fractions of the portion of the spinal cord just ventral to the SDH. aSTEP is significantly decreased.** Saline in lilac, BDNF in dark purple; n = 6. *$p < 0.05$

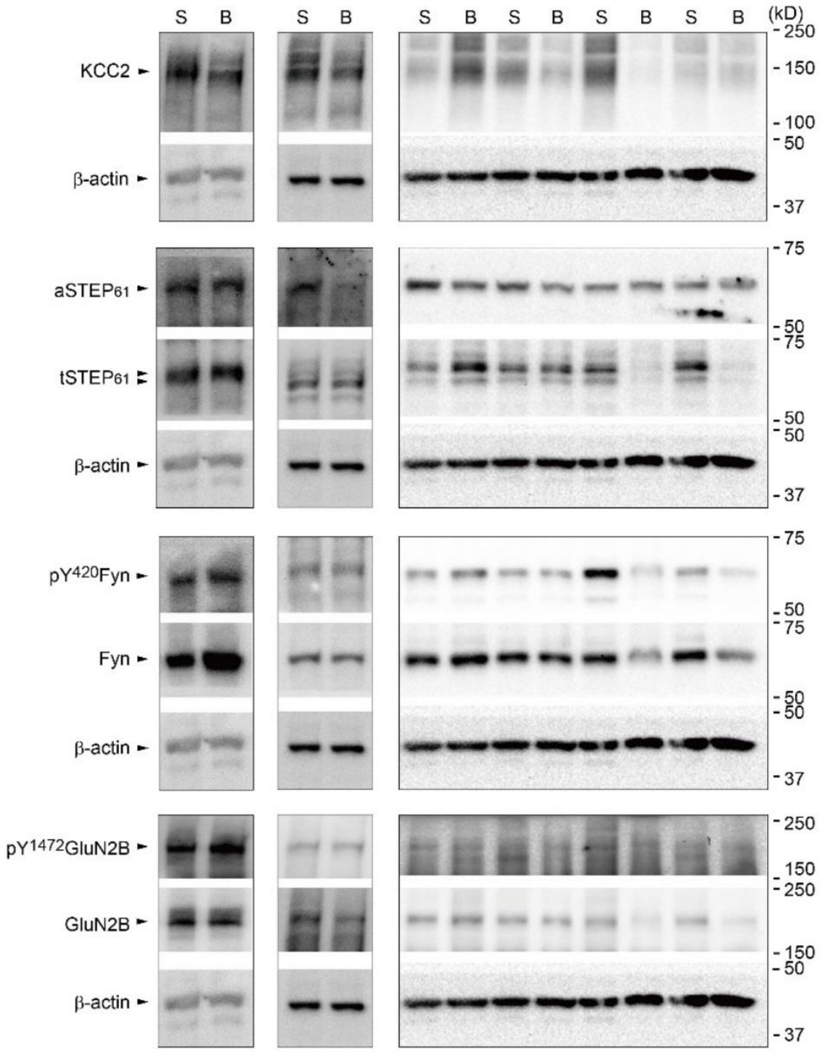


**Supplementary Figure 14. Female BDNF human VH synaptosome gels from human spinal samples treated with either saline (S) or BDNF (B).** Individual gels were cut into sections to allow for probing several targets concurrently β-actin, the loading control, can be seen under each set of targets (labelled on the left while weight of the target, in kD is on the right. n= 6

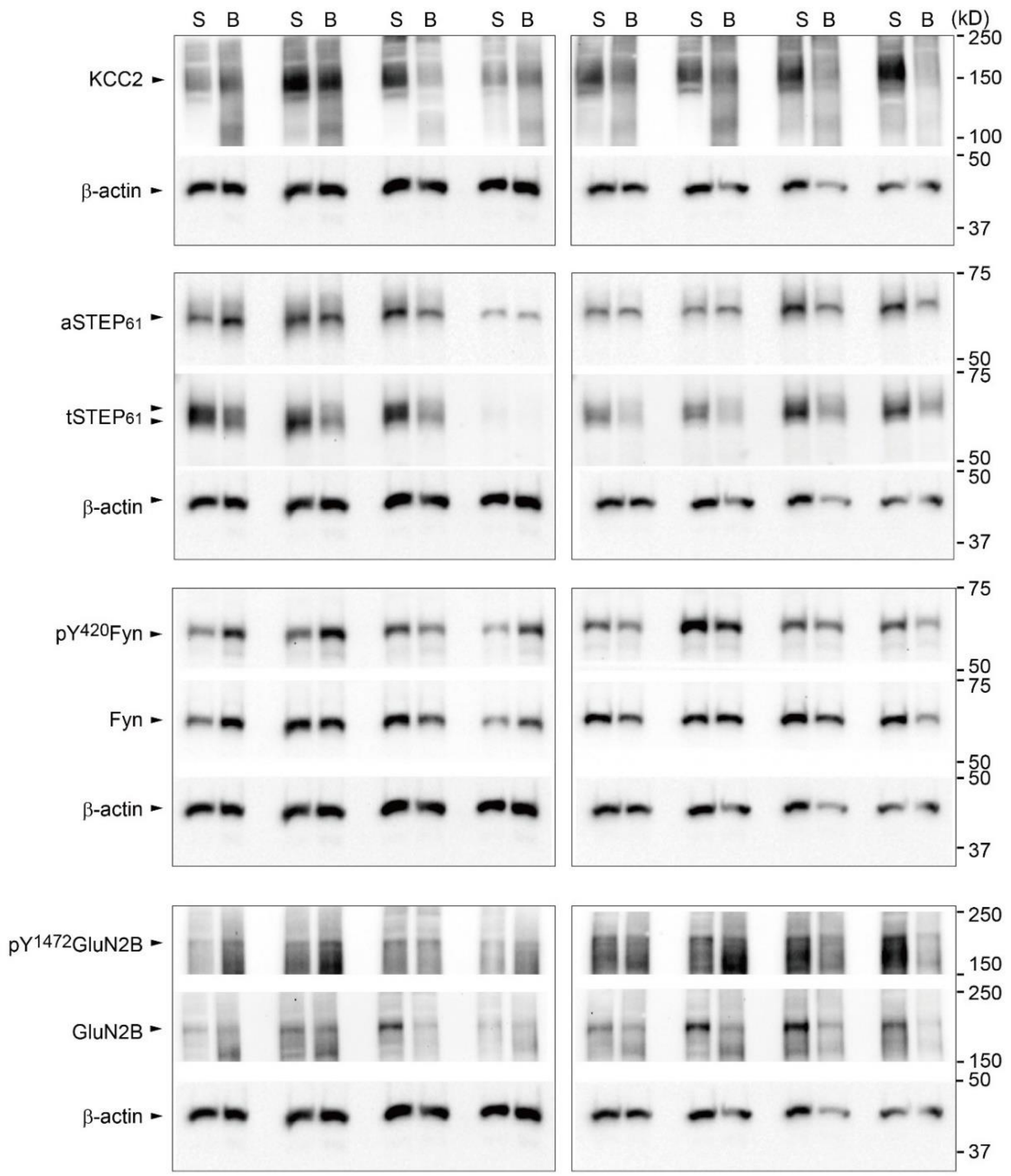


**Supplementary Figure 15. Female OVX rat SDH synaptosome gels from animals treated with either saline (S) or BDNF (B).** Individual gels were cut into sections to allow for probing several targets concurrently β-actin, the loading control, can be seen under each set of targets (labelled on the left while weight of the target, in kD is on the right. n= 8

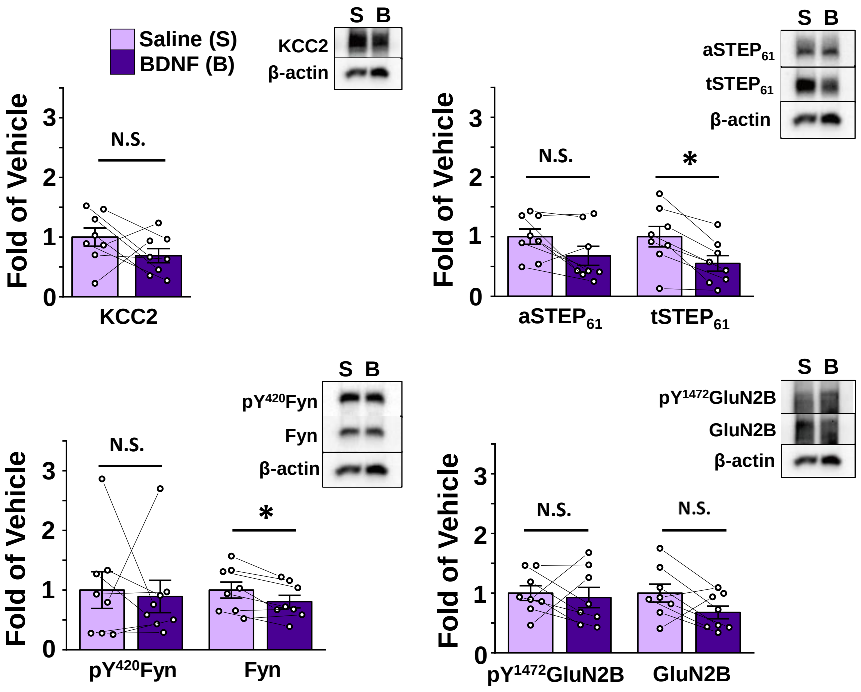


**Supplementary Figure 16. Incubating female OVX rat tissue in 50 ng/mL BDNF elicits decreases in tSTEP and Fyn, and no change in KCC2, aSTEP, pFyn, pGluN2B, and GluN2B in crude synaptosome fractions of the portion of the spinal cord just ventral of the SDH**. Saline in lilac, BDNF in dark purple; n = 8 animals per group.

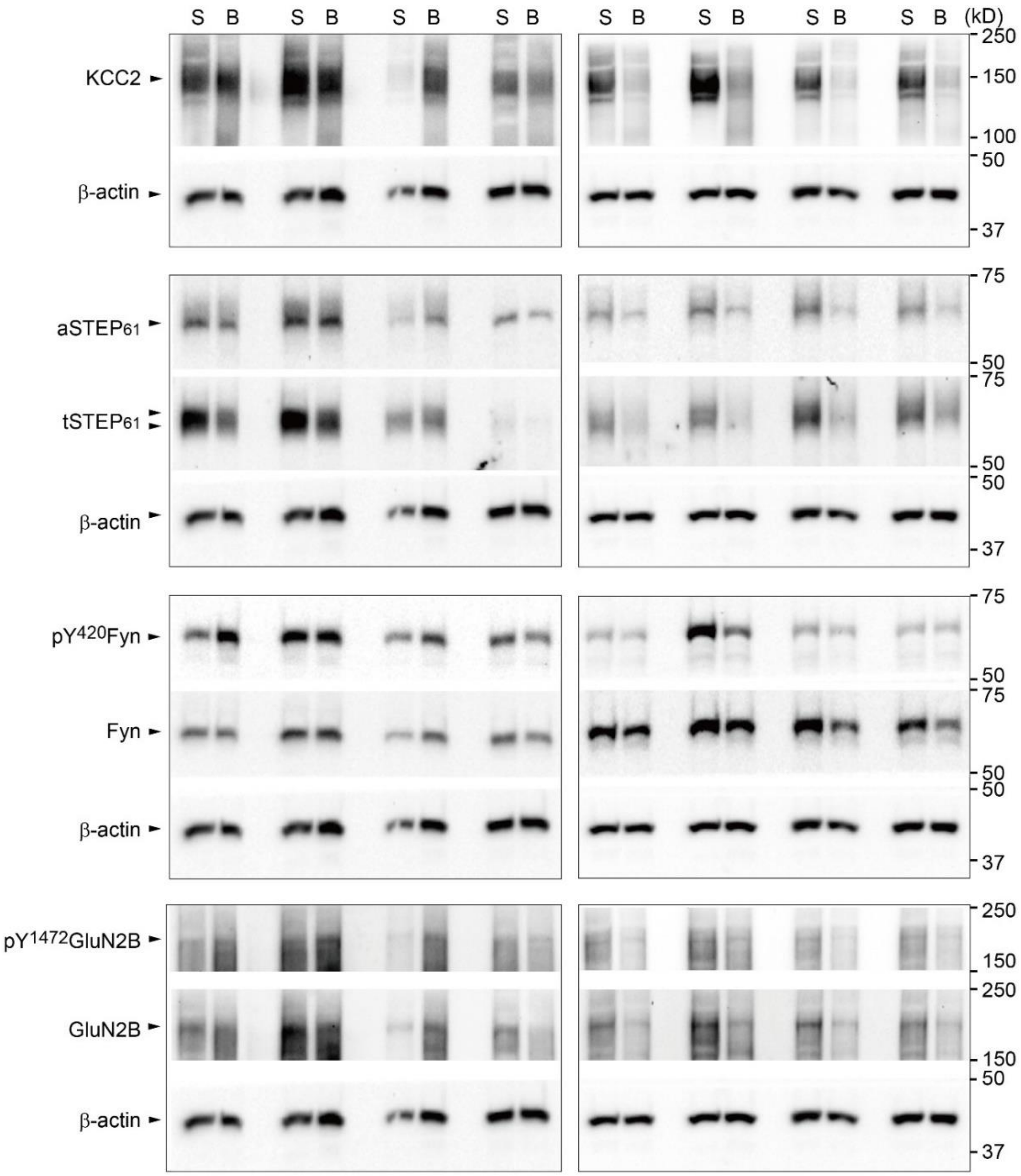


**Supplementary Figure 17. Female OVX rat VH synaptosome gels from animals treated with either saline (S) or BDNF (B).** Individual gels were cut into sections to allow for probing several targets concurrently β-actin, the loading control, can be seen under each set of targets (labelled on the left while weight of the target, in kD is on the right. n= 8

### Supplementary Table 1. Antibodies used for Western Blots.

| Antibody | Format | Immunogen | Host | Target Species | Dilution | Source | Cat # | References |
|---|---|---|---|---|---|---|---|---|
| **Anti-STEP (clone 23E5)** | Monoclonal $IgG_{2b}$ kappa light chain | 18 amino acid sequence mapping at N-terminus of rat STEP46 | Mouse | Mouse, rat | 1:1000 | Santa Cruz Biotechnology | sc-23892 | (Gladding *et al.*, 2014; Jang *et al.*, 2016; Rué *et al.*, 2016; Xu *et al.*, 2017) |
| **Anti-STEP (D9H3)** | Monoclonal IgG | A synthetic peptide corresponding to residues surrounding Ile440 of human STEP61 protein | Rabbit | Mouse, rat, human | 1:1000 | Cell Signaling | 9069S | (Xu *et al.*, 2016) |
| **Anti-KCC2** | Polyclonal | A synthetic peptide mapping at the N-terminus of KCC2 of human origin | Rabbit | Mouse, rat, human | 1:1000 | Santa Cruz Biotechnology | sc-19419-R | (Zhou *et al.*, 2012; Chen *et al.*, 2016) |
| **Anti-Fyn** | Polyclonal | Epitope mapping at the N-terminus of Fyn of human origin | Rabbit | Mouse, rat, human, canine, bovine, porcine, avian | 1:1000 | Santa Cruz Biotechnology | sc-16 | (Brignatz *et al.*, 2009; Levi *et al.*, 2010; Yadav and Denning, 2011) |
| **anti-pY416-Src** | Monoclonal IgG | Produced using synthetic phosphopeptide corresponding to residues surrounding Tyr419 of human Src protein. Detects endogenous levels of Src only when phosphorylated at Tyr416. May cross-react with other Src family members (Lyn, Fyn, Lck, Yes and Hck) when phosphorylated at equivalent sites. May cross-react with overexpressed phosphorylated RTKs. | Rabbit | Human, Mouse, Rat, Monkey | 1:1000 | Cell Signaling | 6943S | (McKinley *et al.*, 2013; Allison *et al.*, 2015; Bieerkehazhi *et al.*, 2017) |
| **Anti-β-actin** | Monoclonal $IgG_1$ kappa light chain | Chicken gizzard actin | Mouse | mouse, rat, human, avian, bovine, canine, porcine, rabbit, Dictyostelium discoideum, Physarum polycephalum | 1:10000 | Santa Cruz Biotechnology | sc-47778 | (Zuo *et al.*, 2009; Ti and Pollard, 2011; Wu *et al.*, 2014) |
| **anti-non-phospho-STEP** | Monoclonal IgG | Produced using synthetic nonphosphopeptide corresponding to residues surrounding Ser221 of human STEP61 protein. | Rabbit | Human, Mouse, Rat | 1:1000 | Cell Signaling Technology | 5659S | (Castonguay *et al.*, 2018) |

| | | | | | | | | |
|---|---|---|---|---|---|---|---|---|
| | | Detects STEP61 protein only when dephosphorylated at Ser221 and of STEP46 protein when dephosphorylated at Ser49 | | | | | | |
| **anti-pY$^{1472}$ GluN2B** | Polyclonal | Affinity Purified from Pooled Serum. Phosphopeptide corresponding to amino acid residues surrounding the phospho-Tyr1472 of NMDA NR2B. | Rabbit | Rat, Mouse, Human, Bovine, Chicken, Non-human primate, Zebra fish, Canine | 1:1000 | PhosphoSolutions | p1516-1472 | (Castillo *et al.*, 2011; Jang *et al.*, 2016) |
| **anti-GluN2B** | Monoclonal IgG | 6His-tagged fusion protein corresponding to amino acids 1265-1464 of mouse NMDA receptor 2A (NR2A) | Rabbit | Mouse, Rat | 1:2000 | Millipore | 06-600 | (Fenster *et al.*, 2012; Wei *et al.*, 2014) |

**Supplementary Table 2. Statistics Summary Table**

| Figure | Comparison | Test | P value | Significant? |
|---|---|---|---|---|
| 1A | PWT – Vehicle | One-way repeated measures ANOVA | Sphericity established: 0.8508 | No |
| 1A | PWT - CFA | One-way repeated measures ANOVA | Sphericity violated, Greenhouse-Geisser: 3.17E-8 | Yes |
| 1A | PWT - CFA: Baseline vs. Day 1 | Pairwise Comp, Bonferroni adj. | 1.4895E-6 | Yes |
| 1A | PWT - CFA: Baseline vs. Day 2 | Pairwise Comp, Bonferroni adj. | 5.1905E-7 | Yes |
| 1A | PWT - CFA: Baseline vs. Day 3 | Pairwise Comp, Bonferroni adj. | 3.5969E-7 | Yes |
| 1B | WB: KCC2 Vehicle vs. CFA | Independent samples t test | 0.05262 | No |
| 1B | WB: aSTEP$_{61}$ Vehicle vs. CFA | Independent samples t test | 0.008495 | Yes |
| 1B | WB: tSTEP$_{61}$ Vehicle vs. CFA | Independent samples t test | 0.5854 | No |
| 1B | WB: pY$^{420}$Fyn Vehicle vs. CFA | Independent samples t test | 0.01325 | Yes |
| 1B | WB: Fyn Vehicle vs. CFA | Independent samples t test | 0.314269 | No |
| 1B | WB: pY$^{1472}$GluN2B Vehicle vs. CFA | Independent samples t test | 5.6060E-4 | Yes |

| | | | | |
|---|---|---|---|---|
| 1B | WB: GluN2B Vehicle vs. CFA | Independent samples t test | 0.02319 | Yes |
| 1C | PWT – Vehicle | One-way repeated measures ANOVA | Sphericity established: 0.1328 | No |
| 1C | PWT - CFA | One-way repeated measures ANOVA | Sphericity violated, Greenhouse-Geisser: 6.2441E-11 | Yes |
| 1C | PWT - CFA: Baseline vs. Day 1 | Pairwise Comp, Bonferroni adj. | 6.1413E-8 | Yes |
| 1C | PWT - CFA: Baseline vs. Day 2 | Pairwise Comp, Bonferroni adj. | 5.7102E-9 | Yes |
| 1C | PWT - CFA: Baseline vs. Day 3 | Pairwise Comp, Bonferroni adj. | 1.2285E-8 | Yes |
| 1D | WB: KCC2 Vehicle vs. CFA | Independent samples t test | 0.7838 | No |
| 1D | WB: $aSTEP_{61}$ Vehicle vs. CFA | Independent samples t test | 0.9580 | No |
| 1D | WB: $tSTEP_{61}$ Vehicle vs. CFA | Independent samples t test, equal variances not assumed | 0.2862 | No |
| 1D | WB: $pY^{420}$Fyn Vehicle vs. CFA | Independent samples t test | 0.1279 | No |
| 1D | WB: Fyn Vehicle vs. CFA | Independent samples t test | 0.8471 | No |
| 1D | WB: $pY^{1472}$GluN2B Vehicle vs. CFA | Independent samples t test | 0.5962 | No |
| 1D | WB: GluN2B Vehicle vs. CFA | Independent samples t test | 0.4250 | No |
| 2A | Male vs. Female Charge Transfer | Independent samples t test | 0.06258 | No |
| 2A | Male vs. Female Decay Constant | Independent samples t test | 0.3221 | No |
| 2A | Male vs. Female Peak Amplitude | Independent samples t test | 0.2183 | No |
| 2B | Male Charge Transfer Control vs. CFA vs. BDNF | Welch's | 1.4040E-5 | Yes |
| 2B | Male Charge Transfer Control vs. CFA | Welch's, Games-Howell | 0.03188 | Yes |
| 2B | Male Charge Transfer Control vs. BDNF | Welch's, Games-Howell | 2.9022E-6 | Yes |
| 2C | Female Charge Transfer Control vs. CFA vs. BDNF | Kruskal Wallis | 0.2540 | No |
| 2D | WB: KCC2 Saline vs. BDNF | Paired samples t test | 0.8796 | No |
| 2D | WB: $aSTEP_{61}$ Saline vs. BDNF | Paired samples t test | 0.7451 | No |
| 2D | WB: $tSTEP_{61}$ Saline vs. BDNF | Paired samples t test | 0.4751 | No |
| 2D | WB: $pY^{420}$Fyn Saline vs. BDNF | Paired samples t test | 0.6427 | No |
| 2D | WB: Fyn Saline vs. BDNF | Paired samples t test | 0.6001 | No |
| 2D | WB: $pY^{1472}$GluN2B Saline vs. BDNF | Paired samples t test | 0.6979 | No |
| 2D | WB: GluN2B Saline vs. BDNF | Paired samples t test | 0.6175 | No |
| 3A | Male Membrane KCC2 Average | Extra sum-of-squares F-test method | 2.0919E-4 | Yes |

| | | | | |
|---|---|---|---|---|
| | Intensity Vehicle vs. BDNF | | | |
| 3A | Male Intracellular KCC2 Average Intensity Vehicle vs. BDNF | Extra sum-of-squares F-test method | 1.7548E-6 | Yes |
| 3B | Female Membrane KCC2 Average Intensity Vehicle vs. BDNF | Extra sum-of-squares F-test method | 0.2511 | No |
| 3B | Female Intracellular KCC2 Average Intensity Vehicle vs. BDNF | Extra sum-of-squares F-test method | 0.7235 | No |
| 3E | Human WB: KCC2 Saline vs. BDNF | Paired samples t test | 0.4691 | No |
| 3E | Human WB: aSTEP$_{61}$ Saline vs. BDNF | Wilcoxon signed-rank test | 0.6002 | No |
| 3E | Human WB: tSTEP$_{61}$ Saline vs. BDNF | Paired samples t test | 0.7730 | No |
| 3E | Human WB: pY$^{420}$Fyn Saline vs. BDNF | Paired samples t test | 0.4945 | No |
| 3E | Human WB: Fyn Saline vs. BDNF | Paired samples t test | 0.6278 | No |
| 3E | Human WB: pY$^{1472}$GluN2B Saline vs. BDNF | Paired samples t test | 0.6002 | No |
| 3E | Human WB: GluN2B Saline vs. BDNF | Paired samples t test | 0.8166 | No |
| 4A | Female: Naive vs. OVX Charge Transfer | Independent samples t test | 0.6602 | No |
| 4A | Female: Naive vs. OVX Decay Constant | Independent samples t test | 0.8513 | No |
| 4A | Female: Naive vs. OVX Peak Amplitude | Mann-Whitney test | 0.9654 | No |
| 4B | Female OVX Charge Transfer ANOVA | One-way ANOVA | 3.5011E-4 | Yes |
| 4B | Female OVX Charge Transfer Control vs. BDNF | One-way ANOVA, Tukey HSD | 5.0100E-4 | Yes |
| 4B | Female OVX Charge Transfer Control vs. BDNF+PP2 | One-way ANOVA, Tukey HSD | .6517 | No |
| 4B | Female OVX Charge Transfer BDNF vs. BDNF+PP2 | One-way ANOVA, Tukey HSD | 0.003389 | Yes |
| 4C | WB: KCC2 Saline vs. BDNF | Paired samples t test | 0.03953 | yes |
| 4C | WB: aSTEP$_{61}$ Saline vs. BDNF | Paired samples t test | 0.4111 | No |
| 4C | WB: tSTEP$_{61}$ Saline vs. BDNF | Paired samples t test | 6.1642E-4 | Yes |
| 4C | WB: pY$^{420}$Fyn Saline vs. BDNF | Paired samples t test | 0.04553 | Yes |
| 4C | WB: Fyn Saline vs. BDNF | Paired samples t test | 0.8685 | No |
| 4C | WB: pY$^{1472}$GluN2B Saline vs. BDNF | Paired samples t test | 0.2441 | No |
| 4C | WB: GluN2B Saline vs. BDNF | Paired samples t test | 0.3252 | No |

| | | | | |
|---|---|---|---|---|
| Supplementary Figure 1 | Male PWT – Vehicle | One-way repeated measures ANOVA | Sphericity violated, Greenhouse-Geisser: 0.7816 | No |
| Supplementary Figure 1 | Male PWT - CFA | One-way repeated measures ANOVA | Sphericity violated, Greenhouse-Geisser: 5.5165E-5 | Yes |
| Supplementary Figure 1 | Male PWT - CFA: Baseline vs. Day 1 | Pairwise Comp, Bonferroni adj. | 2.3591E-3 | Yes |
| Supplementary Figure 1 | Male PWT - CFA: Baseline vs. Day 2 | Pairwise Comp, Bonferroni adj. | 1.7286E-3 | Yes |
| Supplementary Figure 1 | Male PWT - CFA: Baseline vs. Day 3 | Pairwise Comp, Bonferroni adj. | 1.7338E-3 | Yes |
| Supplementary Figure 1 | Male PWT - CFA: Baseline vs. Day 4 | Pairwise Comp, Bonferroni adj. | 1.4770E-3 | Yes |
| Supplementary Figure 1 | Male PWT - CFA: Baseline vs. Day 5 | Pairwise Comp, Bonferroni adj. | 1.2091E-3 | Yes |
| Supplementary Figure 1 | Female PWT – Vehicle | One-way repeated measures ANOVA | Sphericity established: 0.3515 | No |
| Supplementary Figure 1 | Female PWT - CFA | One-way repeated measures ANOVA | Sphericity violated, Greenhouse-Geisser: 1.2645E-7 | Yes |
| Supplementary Figure 1 | Female PWT - CFA: Baseline vs. Day 1 | Pairwise Comp, Bonferroni adj. | 9.3423E-5 | Yes |
| Supplementary Figure 1 | Female PWT - CFA: Baseline vs. Day 2 | Pairwise Comp, Bonferroni adj. | 2.8260E-5 | Yes |
| Supplementary Figure 1 | Female PWT - CFA: Baseline vs. Day 3 | Pairwise Comp, Bonferroni adj. | 9.3423E-5 | Yes |
| Supplementary Figure 1 | Female PWT - CFA: Baseline vs. Day 4 | Pairwise Comp, Bonferroni adj. | 1.1176E-4 | Yes |
| Supplementary Figure 1 | Female PWT - CFA: Baseline vs. Day 5 | Pairwise Comp, Bonferroni adj. | 9.33423E-5 | Yes |
| Supplementary Figure 2 | Male CFA VH: aSTEP$_{61}$ Vehicle vs. CFA | Independent samples t test | 0.9804 | No |
| Supplementary Figure 2 | Male CFA VH: tSTEP$_{61}$ Vehicle vs. CFA | Mann-Whitney test | 0.3823 | No |
| Supplementary Figure 2 | Male CFA VH: pY$^{420}$Fyn Vehicle vs. CFA | Independent samples t test | 0.7686 | No |
| Supplementary Figure 2 | Male CFA VH: Fyn Vehicle vs. CFA | Independent samples t test | 0.9794 | No |
| Supplementary Figure 2 | Male CFA VH: pY$^{1472}$GluN2B Vehicle vs. CFA | Independent samples t test | 0.2592 | No |
| Supplementary Figure 2 | Male CFA VH: GluN2B Vehicle vs. CFA | Independent samples t test | 0.9633 | No |
| Supplementary Figure 5 | Female CFA VH: KCC2 Vehicle vs. CFA | Independent samples t test | 0.7619 | No |
| Supplementary Figure 5 | Female CFA VH: aSTEP$_{61}$ Vehicle vs. CFA | Mann-Whitney test | 1.0000 | No |
| Supplementary Figure 5 | Female CFA VH: tSTEP$_{61}$ Vehicle vs. CFA | Mann-Whitney test | 1.0000 | No |
| Supplementary Figure 5 | Female CFA VH: pY$^{420}$Fyn Vehicle vs. CFA | Independent samples t test | 0.0519 | No |

| | | | | |
|---|---|---|---|---|
| Supplementary Figure 5 | Female CFA VH: Fyn Vehicle vs. CFA | Independent samples t test | 0.1921 | No |
| Supplementary Figure 5 | Female CFA VH: $pY^{1472}$GluN2B Vehicle vs. CFA | Mann-Whitney test | 0.5054 | No |
| Supplementary Figure 5 | Female CFA VH: GluN2B Vehicle vs. CFA | Mann-Whitney test | 0.5737 | No |
| Supplementary Figure 8 | Charge transfer: Female Saline Vs. Long BDNF | Independent samples t test | 0.8880 | No |
| Supplementary Figure 9 | Female VH: KCC2 Saline vs. BDNF | Paired samples t test | 0.8904 | No |
| Supplementary Figure 9 | Female VH: $aSTEP_{61}$ Saline vs. BDNF | Paired samples t test | 0.9093 | No |
| Supplementary Figure 9 | Female VH: $tSTEP_{61}$ Saline vs. BDNF | Paired samples t test | 0.7180 | No |
| Supplementary Figure 9 | Female VH: $pY^{420}$Fyn Saline vs. BDNF | Paired samples t test | 0.7425 | No |
| Supplementary Figure 9 | Female VH: Fyn Saline vs. BDNF | Paired samples t test | 0.3572 | No |
| Supplementary Figure 9 | Female VH: $pY^{1472}$GluN2B Saline vs. BDNF | Paired samples t test | 0.8717 | No |
| Supplementary Figure 9 | Female VH: GluN2B Saline vs. BDNF | Paired samples t test | 0.6093 | No |
| Supplementary Figure 12 | Female VH: KCC2 Saline vs. BDNF | Paired samples t test | 0.5397 | No |
| Supplementary Figure 12 | Female VH: $aSTEP_{61}$ Saline vs. BDNF | Paired samples t test | 0.02462 | Yes |
| Supplementary Figure 12 | Female VH: $tSTEP_{61}$ Saline vs. BDNF | Paired samples t test | 0.4950 | No |
| Supplementary Figure 12 | Female VH: $pY^{420}$Fyn Saline vs. BDNF | Paired samples t test | 0.1819 | No |
| Supplementary Figure 12 | Female VH: Fyn Saline vs. BDNF | Paired samples t test | 0.6616 | No |
| Supplementary Figure 12 | Female VH: $pY^{1472}$GluN2B Saline vs. BDNF | Wilcoxon signed ranks test | 0.2489 | No |
| Supplementary Figure 12 | Female VH: GluN2B Saline vs. BDNF | Paired samples t test | 0.4107 | No |
| Supplementary Figure 15 | OVX Female VH: KCC2 Saline vs. BDNF | Paired samples t test | 0.1779 | No |
| Supplementary Figure 15 | OVX Female VH: $aSTEP_{61}$ Saline vs. BDNF | Wilcoxon signed ranks test | 0.06870 | No |
| Supplementary Figure 15 | OVX Female VH: $tSTEP_{61}$ Saline vs. BDNF | Paired samples t test | 0.001430 | Yes |
| Supplementary Figure 15 | OVX Female VH: $pY^{420}$Fyn Saline vs. BDNF | Wilcoxon signed ranks test | 0.8886 | No |
| Supplementary Figure 15 | OVX Female VH: Fyn Saline vs. BDNF | Paired samples t test | 0.01102 | Yes |
| Supplementary Figure 15 | OVX Female VH: $pY^{1472}$GluN2B Saline vs. BDNF | Paired samples t test | 0.7237 | No |
| Supplementary Figure 15 | OVX Female VH: GluN2B Saline vs. BDNF | Paired samples t test | 0.08205 | No |
